# Scalable human neuronal models of tauopathy producing endogenous seed-competent 4R tau

**DOI:** 10.1101/2025.07.11.664346

**Authors:** Eliona Tsefou, Sumi Bez, Timothy J.Y. Birkle, Martha Foiani, Naoto Watamura, Mathieu Bourdenx, Daria Gavriouchkina, Emir Turkes, Samuel Crawford, Rachel Coneys, Adrian M. Isaacs, Karen Duff

## Abstract

The accumulation of pathological four-repeat (4R) tau is central to several frontotemporal dementia (FTD) subtypes, but human neuronal models amenable to high-throughput screening of 4R tau-targeting therapies remain very limited. To address this, we developed iPSC-derived i^3^Neuron (i^3^N) lines expressing >75% 4R tau, driven by FTD splice-shifting mutations (S305N or S305N/IVS10+3). These neurons develop hyperphosphorylated tau and demonstrate somatodendritic mis-localization. Unlike other stem cell models of tauopathy, these i^3^N neurons develop endogenous seed-competent tau and present pFTAA-positive tau assemblies after 28 days in culture. For scalable screening, we CRISPR-engineered a HiBiT luminescence tag at the endogenous *MAPT* locus into the S305N/IVS10+3 iPSC line, enabling precise quantification of tau levels and pharmacological responses. The model responded predictably to compounds affecting tau clearance, demonstrating its suitability for drug discovery. Overall, this i^3^N platform recapitulates key features of 4R tauopathy and provides a robust system to identify therapeutic modulators of pathological tau.

## Introduction

Tauopathies are a heterogeneous group of neurodegenerative disorders defined by the accumulation of pathological forms of the microtubule-associated protein tau (MAPT) in the brain. These include progressive supranuclear palsy (PSP), corticobasal degeneration (CBD), Pick’s disease, and frontotemporal dementia with Parkinsonism linked to chromosome 17 (FTDP-MAPT) (*1*). In these diseases, tau aggregates into intracellular fibrils that interfere with neuronal function and viability (*2, 3*). Tau exists in six isoforms in the adult human brain, produced by alternative splicing of exons 2, 3, and 10 of the *MAPT* gene (*4*). These isoforms differ in the number of N-terminal inserts and in the presence of either three (3R) or four (4R) microtubule-binding repeats (*5*). During fetal development, the human brain predominantly expresses 3R tau (*6*), but in the healthy adult brain, the ratio of 3R:4R tau is balanced (*7–9*). Disruption of this balance, particularly an excess of 4R tau, is associated with several diseases (*10, 11*).

Human induced pluripotent stem cell (iPSC)-derived neurons are a valuable tool to study early mechanisms of neurodegenerative diseases. However, using such human neuronal models to recapitulate 4R tau expression has historically been difficult, as they typically express predominantly 0N3R tau, reflecting fetal-like tau splicing (*12–16*). Until recently, most of the iPSC-derived neuronal models that were able to induce moderate expression of 4R tau required prolonged and complicated culturing conditions, or they relied on the insertion of multiple tau mutations (*17–21*). Alternative models that make endogenous pathological neuronal tau include directly converted fibroblasts (miNs) (*22*) and 3D organoid cultures from ReNcell VM cells with amyloid co-pathology (*23*).

The S305N mutation, located at the end of exon 10, causes excessive production of 4R tau and consequently leads to FTLD-MAPT (*24–26*). Post-mortem brain tissue from S305N carriers exhibits 4R tau-positive inclusions, including neurofibrillary tangles and ring-like structures surrounding neuronal nuclei, reflecting severe tau pathology (*27, 28*). A recent study has shown that an isogenic set of iPSC lines derived from a S305N mutation carrier has increased 4R tau after three months in culture (*29*). In the current study, we employ i^3^Neuronal (i^3^N) versions of the same isogenic S305N iPSC set. Building on insights from a targeted, humanized mouse model, where combining both S305N and intronic IVS10+3 FTLD-MAPT-related mutations led to the expression of 99% 4R MAPT (*30*). We aimed to determine whether a similar shift could be achieved in a second set of lines derived from a different healthy donor. CRISPR-Cas9-editing was used in the second iPSC line to introduce the S305N point mutation and the IVS10+3 intronic mutation to further enhance 4R levels, as well as a HiBiT tag to monitor tau levels. Elevated 4R tau resulted in tau hyperphosphorylation, changes in the cytoskeleton and, remarkably, the accumulation of endogenous seed-competent tau. We have tested this line with tau-modulating drugs, and we anticipate it can be developed into a platform for drug screening, targeting tau in a range of neurodegenerative diseases.

## Results

### i^3^N neurons with a S305N mutation predominantly express 4R tau after 7 days of differentiation

The S305N mutation affects the RNA stem-loop structure that regulates alternative splicing of exon 10 in the *MAPT* gene (*31*). The IVS10+3 mutation lies adjacent to the splice-donor site and disrupts the stem-loop structure of the pre-mRNA, promoting increased exon 10 inclusion (*32, 33*). To generate our model, we used a commercially available iPSC line from Synthego (802:30F; referred to as WT) and introduced the mutations using CRISPR-Cas9 with a single guide RNA (Fig. S1A). As the IVS10+3 mutation is spliced out, only a single, FTLD-MAPT-causing mutation is present in the protein, and the cells are therefore a valid model of human 4R tauopathy. Two edited, homozygous clones, S305N_C1 and S305N_C2, were isolated and used for experiments. The top 3 off-target effects were validated by PCR and sequencing, but no changes were observed. In addition to these lines, we used a patient-derived S305N heterozygous line (isoHet S305N), along with isogenic wild-type (isoWT) and homozygous (isoHom S305N) lines (Fig. 1A). These lines are available from NCRAD (*29*) and referred to in the current study as 300.12. All iPSC lines were transfected with a doxycycline-inducible human *Neurogenin-2* (hNGN2) transgene using the piggyBac transposon system (Fig. 1A) (*34*). Stable integration of hNGN2 enabled rapid differentiation into human excitatory cortical glutamatergic neurons (i^3^Ns) within three days of doxycycline exposure. Post-transfection, all lines maintained normal karyotypes (assessed via low-coverage whole genome sequencing) and continued to express pluripotency markers (Fig. S1B–E).

**Fig. 1.**
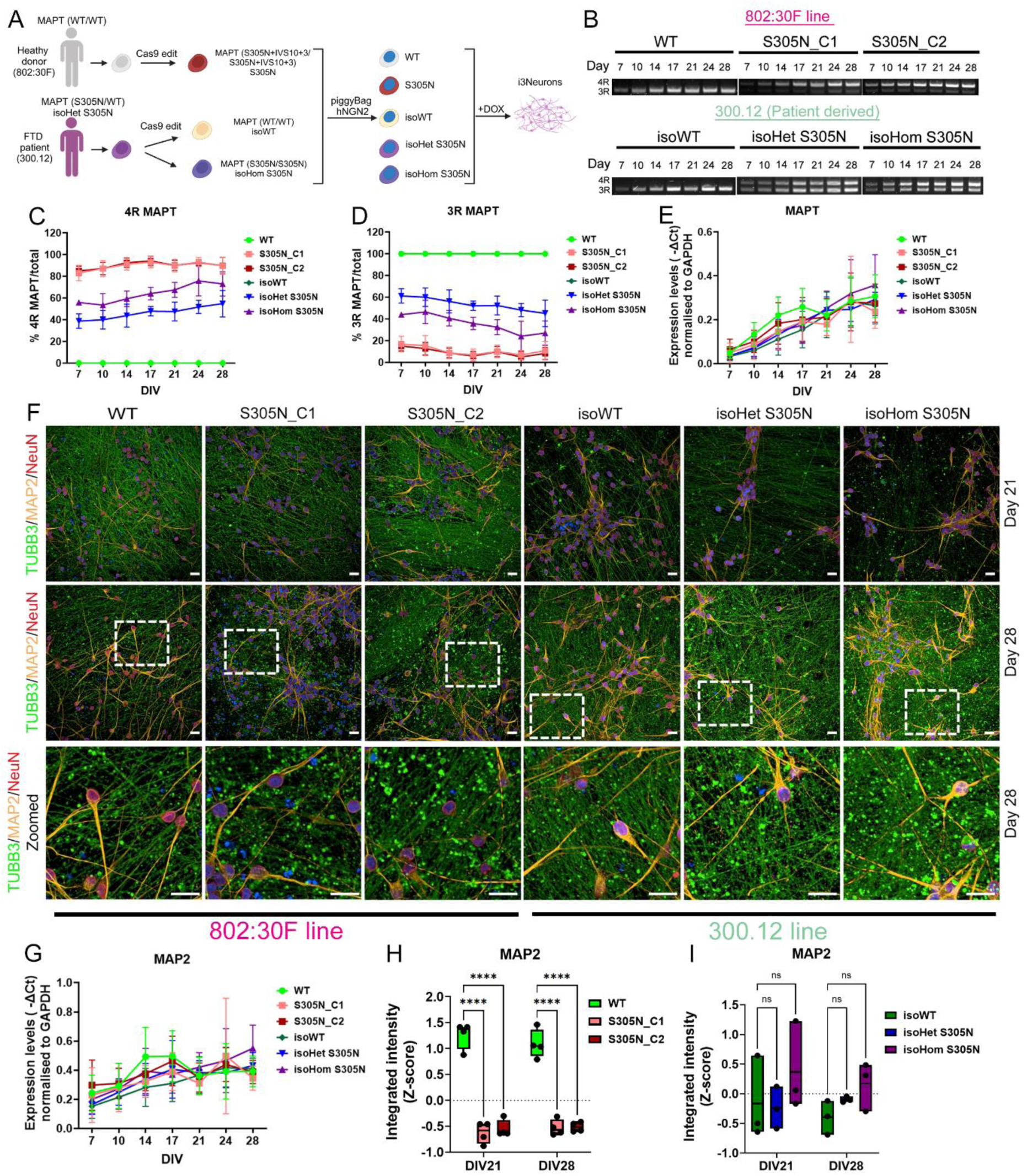
S305N i^3^N neurons express predominantly 4R tau. (A) Schematic representation of the iPSC lines used and the generation of iPSC-derived neurons via the piggyBac hNGN2 system, which enables neuronal differentiation (created with BioRender). (B) Representative semi-quantitative PCR (semi-qPCR) showing 3R and 4R tau isoform levels over time. (C–D) Quantification of 4R tau (C) and 3R tau (D) from panel B (mean ± SD, n = 4–7 independent neuronal differentiations). (E) Total tau mRNA levels measured by qPCR (mean ± SD, n = 4–7 independent neuronal differentiations). (F) Representative images of methanol-fixed i3N neurons at DIV21 and DIV28 stained with TUJ1, MAP2, and NeuN. (G) MAP2 mRNA levels measured by qPCR (mean ± SD, n = 3–5 independent neuronal differentiations). Quantification of MAP2 expression from panel F in the 802:30F (H) and 300.12 (I) iPSC lines (Two-way ANOVA followed by Tukey post-hoc test; ****p < 0.0001; mean ± SD, n = 3–4 independent neuronal differentiations).

To assess the effect of the mutations on exon 10 inclusion, we differentiated all WT and mutant iPSC lines into i^3^N neurons using a two-step protocol adapted from Fernandopulle *et al.* (*35*), culturing them for up to 28 days. We examined MAPT isoform expression by semi-quantitative PCR (semi-qPCR), focusing on the inclusion (4R MAPT) or exclusion (3R MAPT) of exon 10 (Fig. 1B). As expected, both WT and isoWT lines exclusively expressed 3R tau throughout differentiation (DIV7–DIV28). In contrast, both S305N_C1 and C2 clones expressed >80% 4R tau as early as DIV7, increasing to >90% by DIV28. The isoHet S305N line exhibited a 3R:4R ratio of approximately 60:40 at DIV7, shifting to a nearly 50:50 ratio by DIV28. The isoHom S305N line started with 45:55% 3R:4R tau at DIV7, and reached ∼25:75% 3R:4R by DIV28. This line therefore had lower 4R levels compared to the S305N_C1 and C2 clones (Fig. 1C, D). The level of total MAPT mRNA was quantified by RT-qPCR and no change was observed (Fig. 1E). MAPT mRNA could not be detected in the neuronal lines before DIV7.

### Characterization of S305N i^3^N neurons

To assess the effects of the S305N mutation and 4R tau elevation on neuronal maturation, we analysed neuronal marker expression at DIV21 and DIV28 by immunocytochemistry. All i^3^N neuronal lines were stained for βIII-tubulin (TUJ1) (an axonal marker), MAP2 (a somatodendritic marker) and NeuN, a nuclear marker associated with neuronal maturity (Fig. 1F). No difference in TUJ1 expression was observed between S305N_C1 and C2 and the WT control at either timepoint (Fig. S2A). In the isoHet and isoHom S305N lines, TUJ1 expression remained comparable to the isoWT control (Fig. S2B). While overall TUJ1 expression was unchanged, we noted the emergence of irregular, swollen axonal protrusions in all lines expressing the S305N mutation, particularly at DIV28. These structures may represent axonal blebbing, which is commonly associated with neuronal stress, injury, or degeneration (*36, 37*). We quantified the number of blebs in the total TUJ1 area and observed an increase in S305N_C1 and C2, isoHet S305N and isoHom S305N, especially at DIV28, compared to isogenic WT i^3^N neurons (Fig. S2C and D). MAP2 mRNA levels did not differ between lines but increased over time in all neuronal cultures, reaching a plateau after DIV21 (Fig. 1G). However, the level of MAP2 protein was significantly reduced in both S305N_C1 and C2 neurons (Fig. 1H), whereas no change was detected in the isoHet and isoHom S305N lines (Fig. 1I). The observed reduction in MAP2 protein in the S305N_C1 and C2 lines may point to cytoskeletal instability, which coincides with the blebbing in these neurons. NeuN staining revealed that over 80% of cells in all lines were NeuN-positive, indicating a high proportion of mature neurons (Fig. S1E).

We also examined the expression of VGLUT1, a marker of glutamatergic neurons. VGLUT1 mRNA levels decreased over time, reaching a plateau around DIV17 (Fig. S2F). At the protein level, immunocytochemistry at DIV21 and DIV28 revealed a statistically significant reduction in VGLUT1 expression in the S305N_C1 and C2 lines, with C1 showing nearly a 50% decrease compared to WT (Fig S2G and H). Both S305N_C1 and C2 presented similar changes; however, for some measurements, a stronger phenotype was evident in the S305N_C1 neurons. No difference was observed in the isoHet and isoHom S305N lines at the timepoints studied (Fig. S2I), which may reflect the lower levels of 4R (or higher relative levels of 3R tau) in these lines.

### Mis-localization and distribution of tau in neurons expressing S305N mutant isoforms

Tau undergoes proteolytic cleavage by various proteases, generating fragments that contribute to the pathogenesis of several neurodegenerative disorders (*38, 39*). To investigate the subcellular distribution of tau, we performed immunocytochemistry on methanol-fixed neurons at DIV21 and DIV28 using antibodies targeting distinct tau domains: Tau13 (N-terminus, epitope 2–18 aa) and TauC (C-terminus, epitope 244–441 aa; Fig. 2A and Fig. S3A for S305N_C2). In both WT and isoWT neurons, Tau13 and TauC labelled tau both in the somatodendritic compartment (co-localized with MAP2, Fig. 2B) and in the axons (Fig. S3B, co-localized with TUJ1), with signal intensity increasing between DIV21 and DIV28 (Fig. 2B). S305N_C1 and C2 neurons displayed pronounced somatodendritic tau retention with markedly reduced axonal staining (Fig. 2B and Fig. S3A and B). Densitometric analysis revealed an >80% reduction in total tau signal detected by Tau13 and TauC antibodies in this group (Figure 2C and E), with both clones presenting similar tau reduction. In the isoHet and isoHom S305N mutant neurons, tau remained detectable in both the somatodendritic compartment and axons (Fig. 2B and Fig. S3B). Among the antibodies, Tau13 showed the least variability and revealed significantly reduced tau levels in isoHet neurons at DIV28, with a trend toward reduction observed in isoHom neurons (Fig. 2D and F, 22% reduction, p = 0.0894).

**Fig. 2.**
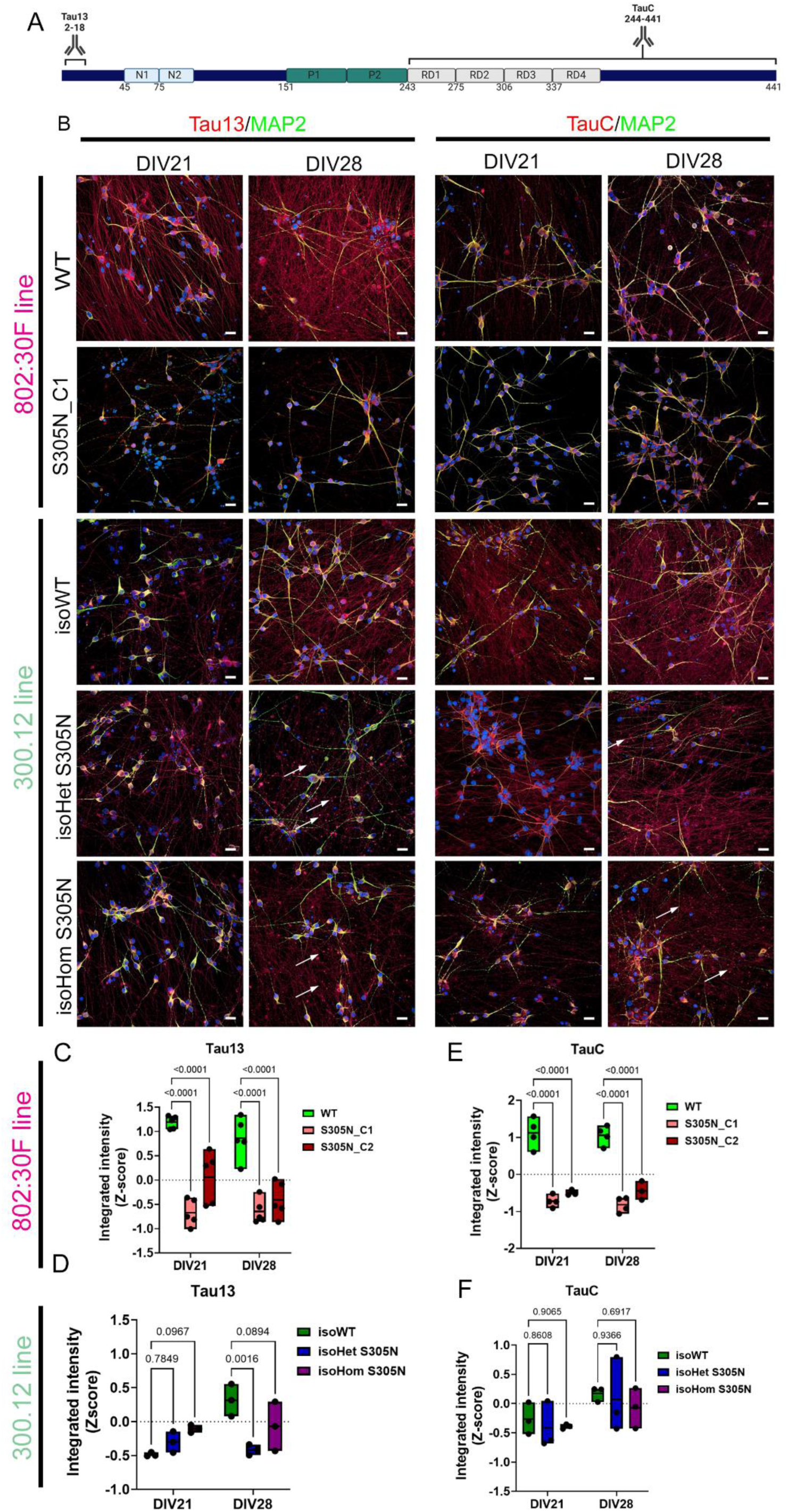
Distribution of total tau in S305N i^3^N neurons. (A) Schematic map of full-length tau and the epitope sites targeted by the total tau antibodies used (created with BioRender). (B) Representative images of DIV21 and DIV28 i3N neurons double-stained with MAP2 and one of the total tau antibodies: Tau13 (left) and TauC (right). (C–D) Quantification of Tau13 staining, and (E–F) quantification of TauC staining across all neuronal lines compared to their isogenic wild-type (WT) controls. (C–F) Data analyzed by two-way ANOVA followed by Tukey post-hoc test; mean ± SD, n = 3–5 independent neuronal differentiations.

Furthermore, neurons expressing isoHet and isoHom S305N tau displayed distinct tau-positive “blebs” at DIV28 when stained with Tau13 and TauC (Fig. 2B, arrows). Co-staining with TUJ1 revealed partial colocalization of these tau “blebs” with tubulin-rich structures (Fig. S3B, see arrows). Overall, these results suggest that high levels of 4R tau relative to 3R (or the effect of homozygous S305N mutation) led to both mis-localization and altered levels of tau, particularly affecting its axonal distribution. The most profound changes were observed in S305N_C1 and C2 neurons, where tau was almost exclusively confined to the somatodendritic compartment. The fact that both C1 and C2 showed a similar effect on total tau indicates that the observed results are not due to clonal selectivity or because of random integration of the hNGN2 cassette. While tau remained detectable in axons of isoHet and isoHom neurons, overall levels compared to isoWT trended lower, especially as revealed by Tau13 and TauC staining.

### Increased tau phosphorylation in neurons expressing S305N mutant isoforms

To investigate the effects of the S305N mutation on tau phosphorylation, we examined the relative levels and distribution of phosphorylated tau (pTau) epitopes using the following antibodies: AT8 (pSer202/pThr205), CP13 (pSer202) and AT100 (pThr212/pSer214). Immunocytochemistry was performed on methanol-fixed neurons at DIV21 and DIV28 (Fig. 3A and Fig. S4A for S305N_C2). In S305N_C1 and C2 neurons, AT8 staining was predominantly localized to the somatodendritic compartment, whereas in the WT neurons AT8 staining was much lower (Fig. 3B). Quantitative analysis revealed a significant increase in AT8 signal at DIV21 in the S305N_C2 only (p = 0.025, whereas at DIV28 AT8 levels did not change (Fig. 3C). A similar trend was observed for S305N_C1, but that change did not reach significance. When normalizing AT8 levels to total tau (TauC) we observed a significant increase in the S305N_C1 only, at both timepoints (Fig. S4C; AT8/TauC staining not shown). CP13 staining in S305N_C1 and C2 neurons showed both somatodendritic and axonal localization. However, total CP13 signal was significantly reduced compared to WT (Fig. 3D). After normalization to TauC, a trend toward increased CP13 levels was observed at DIV28 (Fig. S4B and D). Surprisingly, AT100 staining in S305N_C1 and C2 neurons was restricted exclusively to the axons (Figure 3B). Given the overall reduction in total tau in this line, the presence of detectable AT100 signal suggests this form of tau was still localized within the axonal compartment. Interestingly, AT100 diverged between genotypes: while signal decreased over time in WT neurons, it progressively increased in S305N_C1 and C2 neurons (Fig. 3E).

**Fig. 3.**
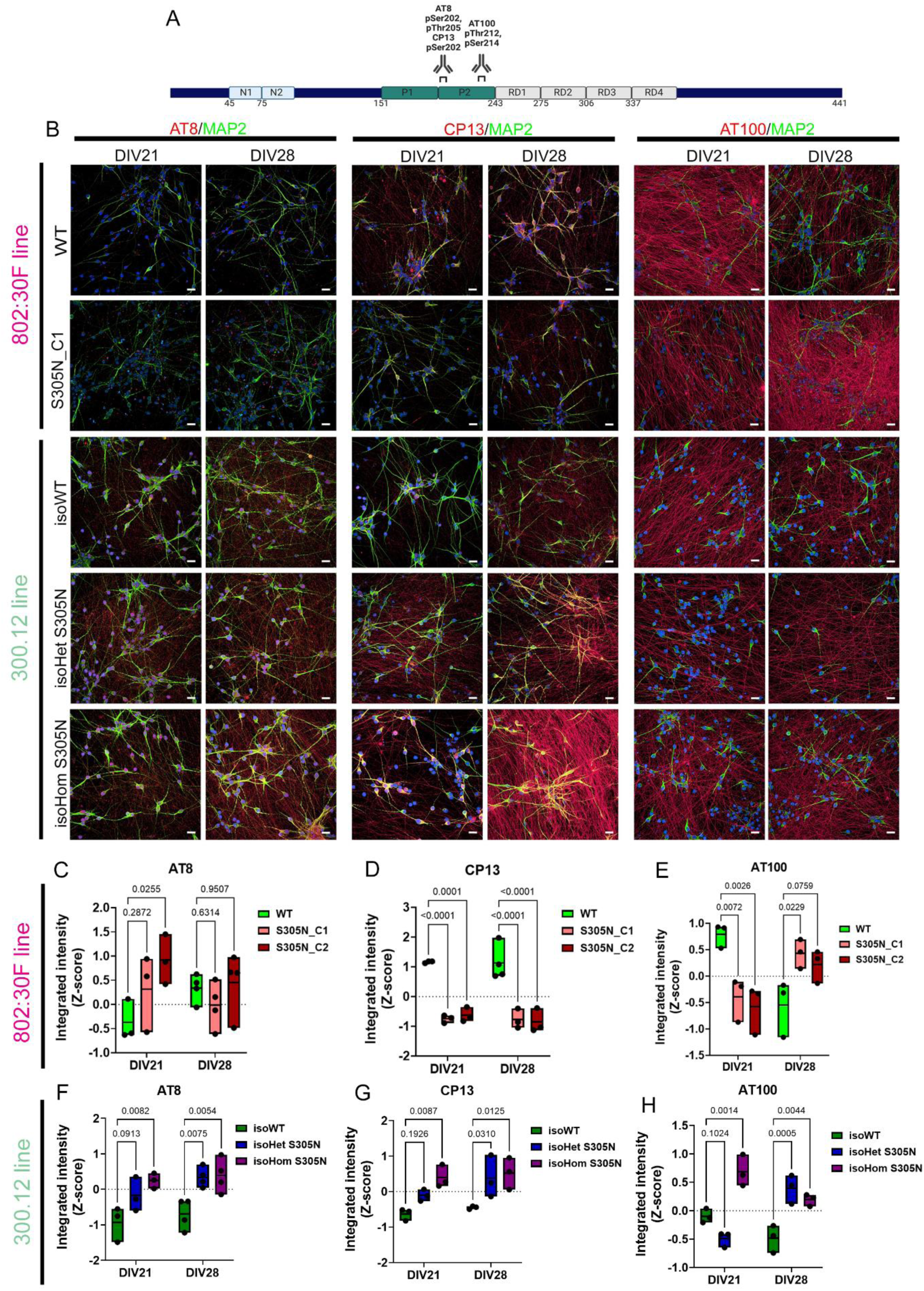
Distribution of phosphorylated tau in S305N i^3^N neurons. (A) Schematic map of full-length tau and the epitope sites recognized by the phosphorylated tau antibodies used (created with BioRender). (B) Representative images of DIV21 and DIV28 i3N neurons double-labelled with MAP2 and one of the phosphorylated tau antibodies: AT8 (left), CP13 (middle), and AT100 (right). (C–H) Quantification of AT8 (C–D), CP13 (E–F), and AT100 (G–H) staining across all neuronal lines compared to their isogenic wild-type (WT) controls. Statistical analysis was performed using two-way ANOVA followed by Tukey post-hoc test; data are presented as mean ± SD, n = 3–4 independent neuronal differentiations.

In isoHet and isoHom S305N neurons, AT8 and CP13 signal was evident in both soma and axons. Levels increased in a dose-dependent manner relative to S305N allele copy number compared to their isogenic WT controls at DIV21 (Fig. 3B, F, G). At DIV28, both AT8 and CP13 levels were significantly elevated in isoHet and isoHom neurons. AT100 in the isoWT neurons mirrored the trend seen in WT neurons, showing a reduction over time. In isoHet neurons, the level of AT100 significantly increased at DIV28 (p=0.0005), whereas in isoHom neurons, elevated AT100 signal was detected at both timepoints (Fig. 3H). Notably, unlike AT8 and CP13, AT100 did not exhibit a clear allele-dose-dependent increase.

Collectively, these results highlight distinct phosphorylation patterns and subcellular localization profiles of tau in neurons with elevated 4R tau, expressing the S305N mutation. Some phospho-epitopes, such as AT8, were restricted to the soma (e.g., in S305N_C1 and C2), others were distributed in both soma and axons (e.g., CP13), while AT100 was found exclusively in axons. Increased 4R tau in S305N mutant neurons therefore influenced isoform composition, the phosphorylation status and the spatial distribution of tau proteoforms.

### Temporal analysis of total and phosphorylated tau reveals early alterations in S305N mutant neurons

To determine whether the observed reduction in total tau levels in S305N_C1 neurons at DIV21 and DIV28 represents a developmental event or a progressive consequence of differentiation, we performed immunoblotting on lysates collected from DIV7 to DIV28. Blots were probed with antibodies against total tau (Tau13 and TauC) and phosphorylated tau (pTau) (CP13 and AT8; Fig. 4A). Across all timepoints, S305N_C1 and C2 neurons displayed consistently lower levels of total tau compared to WT controls (Fig. S5A and B). Tau protein expression in both WT and S305N_C1 and C2 neurons increased over time, mirroring MAPT mRNA expression patterns. As previous studies focused on DIV21 and DIV28, we specifically compared tau expression at these timepoints, as well as at DIV10, the earliest timepoint where tau was detected, with all antibodies used (Fig. 4B, and C). In the isoHet S305N neurons, Tau13, was significantly increased at DIV10 (Fig. 4B left graphs), whereas at DIV21 and DIV28, Tau13 levels presented a trend toward reduction compared to isoWT (p=0.06; Fig. 4B middle and right panel). In the isoHom S305N neurons, a ∼40% reduction in Tau13 was observed at DIV21 and DIV28, while TauC showed a trend toward reduced levels (∼20% reduction, Fig. C). Notably, Tau13 and TauC changes mirrored immunocytochemistry results. A trend towards reduced total tau levels in the same lines was also observed in a previous study when using the direct differentiation approach to generate cortical neurons (*29*). Generally, the reduction in total tau levels was seen in neurons expressing more than 50% 4R tau.

**Fig. 4.**
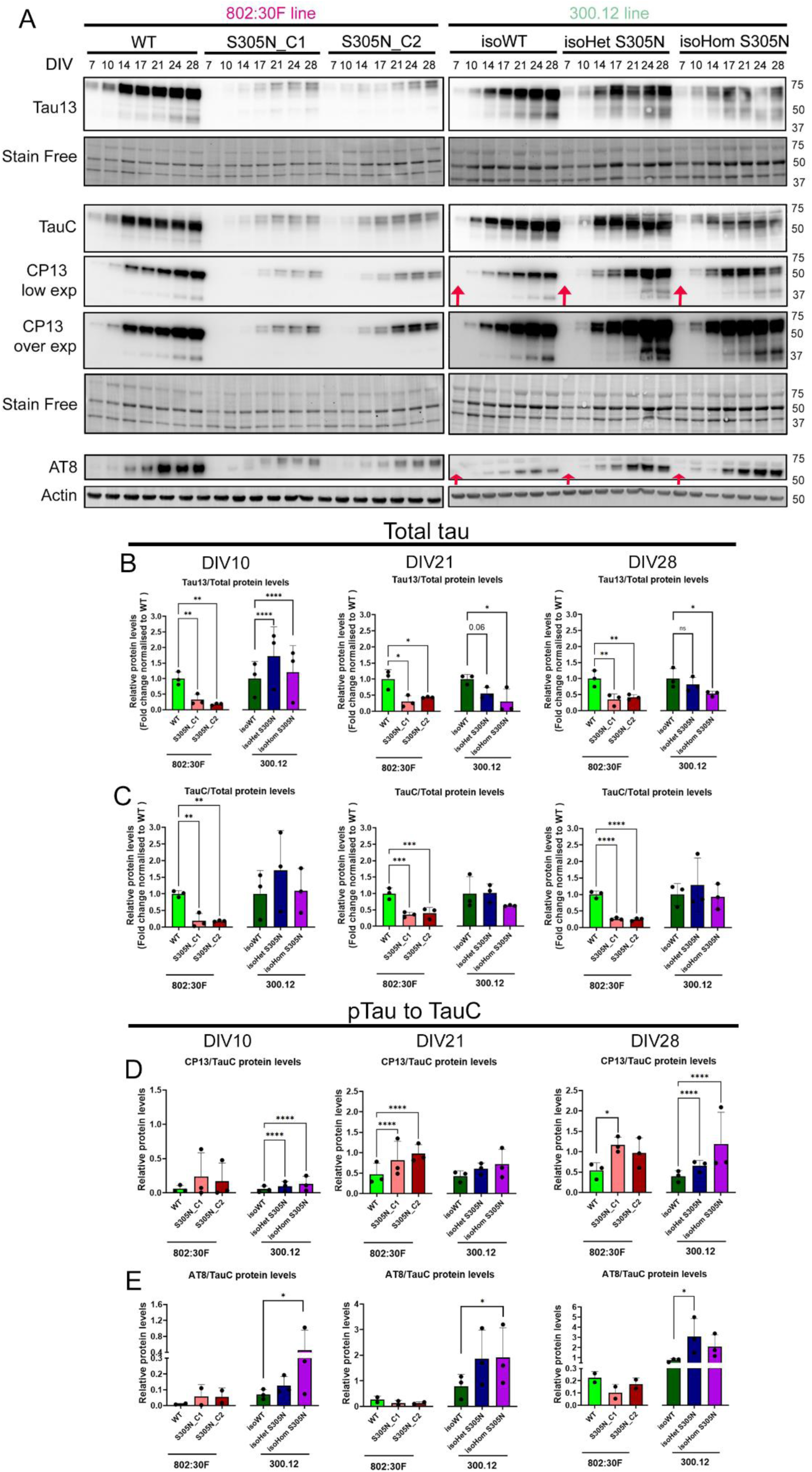
Temporal analysis of total and phosphorylated tau in S305N i^3^N neurons. (A) Representative immunoblots showing total tau (Tau13: top; TauC: bottom) and phosphorylated tau (pTau: CP13 and AT8, bottom) in all neuronal lines over time. Stain-free imaging or actin was used as a loading control, as indicated. Red arrows denote early phosphorylation events. (B–C) Quantification of total tau antibody levels: Tau13/total protein (B) and TauC/total protein (C) at DIV10, DIV21, and DIV28 across all lines. (D–E) Quantification of phosphorylated tau antibody levels: CP13/TauC (D) and AT8/TauC (E) at DIV10, DIV21, and DIV28 across all neuronal lines. Statistical comparisons were made between mutant lines and their respective isogenic wild-type (WT) controls. Repeated measures (RM) one-way ANOVA with Dunnett’s post-hoc test was used to compare S305N_C1 and C2 vs WT, and isoHet/isoHom S305N vs isoWT. Immunoblots for each line were performed separately. These tests were selected due to variability in immunoblot replicates. Data are presented as mean ± SD; *p < 0.05, **p < 0.01, ***p < 0.001, ****p < 0.0001; n = 2–3 independent neuronal differentiations

Phosphorylated tau levels showed a similar pattern to tau in the S305N_C1 and C2 neurons over time (Fig. S5C-F). CP13 and AT8 expression were significantly reduced in S305N_C1 and C2 neurons when normalized to total protein at DIV10, 21 and 28 (Fig. S5G-I). When normalized to TauC, the reduction in CP13 was attenuated and a trend toward increased levels was observed at DIV10, which became statistically significant at DIV21 and DIV28 in the S305N_C1 and C2 compared to WT neurons (Fig. 4D). Interestingly, the ratio of AT8/TauC was elevated at earlier timepoints (DIV10), but declined thereafter, suggesting an early burst of phosphorylation at pSer202/pThr205 (Fig. 4E). It is important to note that, based on immunocytochemistry, AT8/TauC levels were increased, possibly due to differences in detecting phosphorylated epitopes between techniques, which can affect accessibility. Elevated CP13 levels normalized to total tau indicate tau hyperphosphorylation in S305N_C1 and C2 neurons.

Consistent with staining data, both CP13 and AT8 levels were elevated in isoHet and isoHom neurons over time (Fig. S5C-F). When normalized to total protein, CP13 expression increased in the isoHet S305N relative to isogenic WT controls at DIV21and DIV28 only (Fig. S5G-I; graphs on the left). In the isoHom S305N, CP13 levels were also increased over time but did not reach significance due to variability (Fig. S5G-I; graphs on the left). AT8 levels followed a similar pattern but did not reach significance at any timepoint due to variability (Fig. S5G-I; graphs on the right). Both CP13 and AT8 were also normalised to total tau. CP13/TauC was statistically significantly increased in isoHom and isoHet S305N neurons at DIV10 and DIV28 and presented a trend towards increase expression at DIV21 (Fig. 4D). Similarly, when we normalized AT8 levels to TauC a significant increase in AT8 levels was observed in the isoHom S305N compared to isoWT neurons at DIV10 and DIV21 (Fig. 4E). AT8/TauC levels were increased in the isoHet S305N, but that change was only significant at DIV28 (Fig. 4E). A 4R MAPT-dependent increase in AT8/TauC levels was also noted at DIV10 and for CP13/TauC at DIV28 (Fig. 4E,). The increase in CP13/TauC and AT8/TauC indicated that tau is hyperphosphorylated in the isoHet and isoHom S305N neurons. Notably, both AT8 and CP13 signals were detectable as early as DIV7 in both neuronal lines, indicating early tau phosphorylation (Fig. 4A red arrows and Fig. S5J).

To further characterize the isoform composition of tau in each neuronal genotype, protein samples were dephosphorylated and analysed by Western blot with Tau13 immunolabelling. As expected, WT and isoWT neurons expressed predominantly 0N3R tau at DIV21 and DIV28, consistent with tau splicing profiles in iPSC-derived neurons (Fig. S5K). In contrast, S305N_C1 neurons expressed only 0N4R tau across both timepoints. IsoHet neurons exhibited a 1:1 ratio of 0N3R to 0N4R tau, while isoHom neurons expressed 58% 0N4R tau at DIV28.

These findings confirm that S305N_C1 and C2 neurons consistently express only 4R tau during differentiation. Although isoHet and isoHom neurons also showed reduced total tau after DIV21, both lines exhibited substantial phosphorylated tau accumulation early in development. This early, and 4R MAPT-dependent phosphorylation of tau, particularly in the context of elevated 4R tau, may underlie the pathophysiological changes associated with mutations that increase exon 10 inclusion, as seen in post-mortem human tissue (*25, 27, 28, 40, 41*).

### S305N mutant neurons develop endogenous tau oligomers and seeds

To evaluate the aggregation potential of tau in S305N mutant neurons, we performed immunocytochemistry using the tau oligomer-recognizing antibody TOC1 developed by Lester Binder’s lab (*42*), on methanol-fixed cells at DIV21 and DIV28. Neurons were co-labelled with MAP2 and TUJ1 to assess subcellular localisation. At DIV28, TOC1 signal was undetectable in WT and isoWT neurons; in contrast, neurons harbouring the S305N mutation exhibited markedly increased TOC1 labelling (Fig. 5A and Fig. S6C for S305N_C2). In S305N_C1 and C2 neurons, TOC1 signal was predominantly perinuclear and strongly colocalised with MAP2 (see enlarged MAP2/TOC1 panels). Notably, there was minimal colocalization with TUJ1 or tubulin “blebs” in these neurons. Both isoHet and isoHom S305N neurons also displayed elevated TOC1 labelling at DIV28, with isoHet neurons appearing to exhibit slightly higher levels. In these cells, TOC1 occupied a significant portion of the MAP2-positive area, with occasional puncta extending into TUJ1-positive regions. In some cases, these puncta colocalised with tubulin blebs (Fig. 5A TUJ1/MAP2/TOC1 panels, arrows). Quantitative analysis confirmed a significant increase in TOC1 signal in all S305N-mutant neurons compared to their isogenic WT controls at DIV28 (Fig. 5B). To confirm that TOC1-positive puncta reflected accumulated tau, we performed co-staining with Tau13. Colocalization between TOC1 and Tau13 was observed in DIV28 neurons, supporting the specificity of TOC1 for tau species (Fig. S6C). At DIV21, TOC1 signal was also detectable and largely restricted to MAP2-positive somatodendritic compartments (Fig. S6A). Colocalization of TOC1 with tubulin blebs was again observed in isoHom neurons (Fig. S6B). Statistically significant increases in TOC1 signal were found in S305N_C1 and C2 and isoHet neurons at this earlier timepoint (Fig. S7A). We also confirmed that the observed phenotype is not driven solely by a few TOC1-positive i3N neurons. To support this, we calculated the percentage of neurons with positive TOC1 staining within the MAP2-defined area, and found that at both time points, all mutant lines exhibited over 60% TOC1-positive staining (Fig. S7B).

**Fig. 5.**
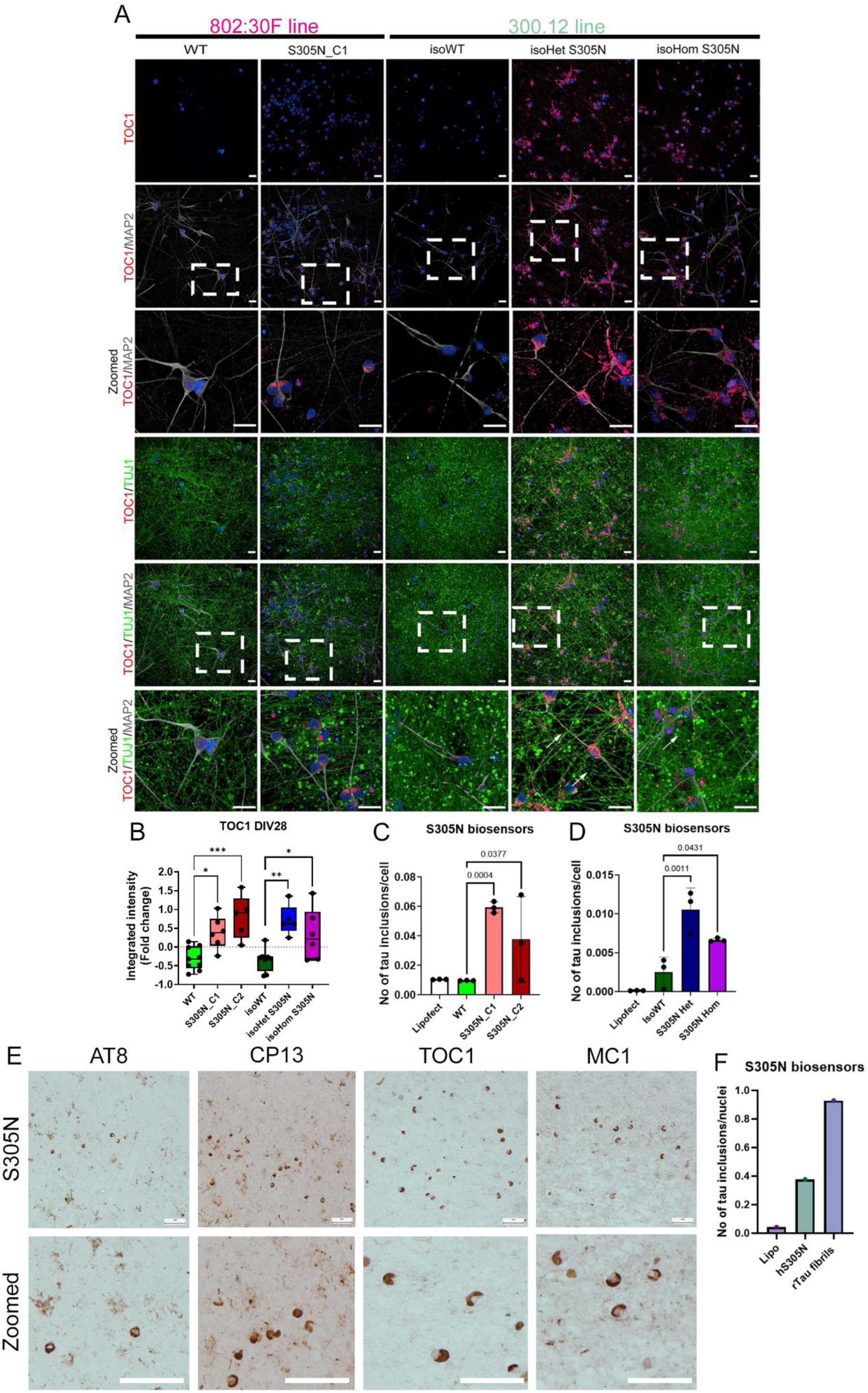
S305N i^3^N neurons form endogenous seed-competent tau. (A) Representative images of DIV28 i^3^N neurons co-labelled with TOC1 and either MAP2 (top panel and zoomed-in) or TUJ1 (bottom panel and zoomed-in) across all neuronal lines. (B) Quantification of TOC1 labelling in DIV28 i3N neurons. Statistical analysis was performed using one-way ANOVA followed by Tukey post-hoc test to compare mutant lines with their respective isogenic controls (mean ± SD, n = 4–5 independent neuronal differentiations). (C–D) Quantification of seeding activity at DIV28 in S305N biosensor lines (one-way ANOVA followed by Tukey post-hoc test; mean ± SD, n = 3 independent neuronal differentiations). (E) Representative images of human brain tissue from patients carrying the S305N mutation, stained for AT8, CP13, TOC1, and MC1. (F) Quantification of seeding activity from 3 µg of frozen human S305N brain tissue using the S305N biosensor assay. Statistical significance: p < 0.05, **p < 0.01, ***p < 0.001.

To assess whether S305N neurons developed tau seeds, we performed both a FRET-based seeding assay and a modified imaging seeding assay (*43*) using a tau biosensor cell line expressing either the repeat domain of tau with the S305N mutation fused to YFP (S305N biosensor (*30*)) or the P301S mutation (P301S biosensors (*44*)). 18 µg of TBS-soluble cell extracts from DIV28 S305N_C1 and C2, isoHet, and isoHom neurons triggered a significant increase in tau inclusion formation compared to their respective isogenic WT controls (Fig. 5C, D, Fig. S7C) in the S305N biosensor. With the P301S biosensor, we were also able to detect increased FRET signal in all lines compared to isogenic WT, however, that change was only significant in the S305N_C1 line, probably due to variation between inductions (Fig. S7D and E). In contrast, extracts from DIV21 neurons did not induce seeding activity (Fig. S7 F and G), suggesting that seed-competent tau develops later in neuronal maturation. Collectively, these results indicate that S305N mutant neurons develop robust TOC1-positive assemblies and seed-competent tau species capable of propagating aggregation in a cellular biosensor model.

Post-mortem human frontal cortex brain tissue from an S305N mutation carrier demonstrated tau pathology consistent with our findings in i^3^N neurons. Neurons in the human S305N case were positive for AT8, CP13, TOC1 and MC1 (Fig. 5E). Ring-shaped tau aggregates surrounding neuronal nuclei were visualised using TOC1 and MC1 antibodies. These observations are consistent with previously reported pathological features identified in human S305N FTLD-MAPT tissue (*25, 27, 28, 30, 40, 41*). 3 µg of TBS-soluble extract from S305N human brain tissue was able to template the aggregation of tau in the S305N biosensor line (Fig. 5F and Fig. S7H) and the P301S line (Fig. S7I). Overall, our S305N i^3^N neurons recapitulate key aspects of human pathology except for MC1 immunoreactivity. Given the relatively short culture times, these cells may therefore reflect the earliest tau-related pathological changes observed in humans.

### Live imaging reveals progressive accumulation of tau assemblies in S305N neurons

Seeding assays indicated that neurons carrying the S305N mutation develop seed-competent tau by DIV28, but not at DIV21. Given that TOC1-positive signal was detected at both timepoints, we next sought to monitor the longitudinal formation and progression of tau assemblies in live neuronal cultures. To accomplish this, we used the conformation-sensitive fluorescent dye pentameric formyl thiophene acetic acid (pFTAA), which selectively binds to β-sheet-rich protein aggregates, including both early tau oligomers and mature fibrils (*45, 46*). Previous studies have shown that pFTAA is non-toxic and remains stable in primary neuronal cultures for extended periods with routine media replacement (*46*). pFTAA and a far-red live tubulin dye were applied to neurons at DIV25, and the accumulation of tau assemblies was imaged over a 72-hour period (Fig. 6A). In S305N_C1 and C2 neurons, pFTAA-positive puncta were first observed within axonal processes, identified by tubulin staining. Over time, larger aggregates formed, predominantly in pre-axonal segments or in proximity to the nucleus (arrows in enlarged panel, Fig. 6A and Fig. S8A for S305N_C2). Quantification revealed that pFTAA-positive puncta began to emerge approximately 18 hours after dye application and progressively increased over time in the S305N_C1 and C2 neurons (Fig. 6B). Similarly, isoHet and isoHom S305N neurons also showed pFTAA accumulation, but as early as 30 minutes after incubation (DIV25), indicating a more rapid onset of aggregate formation or that aggregates are already present at the starting timepoint (Fig. 6A). Some pFTAA-positive structures appeared in regions formerly occupied by viable neurons which could indicate the structures were associated with neuronal death, while others extended from pre-existing pFTAA accumulations. These puncta were consistently observed in tubulin-rich regions and along axon-like projections (Fig. 6A, arrows). Quantitative analysis confirmed a steady increase in pFTAA signal in both isoHet and isoHom S305N neurons throughout the imaging period (Fig. 6C). WT and isoWT neurons, in contrast, exhibited very little pFTAA signal at all timepoints.

**Fig. 6.**
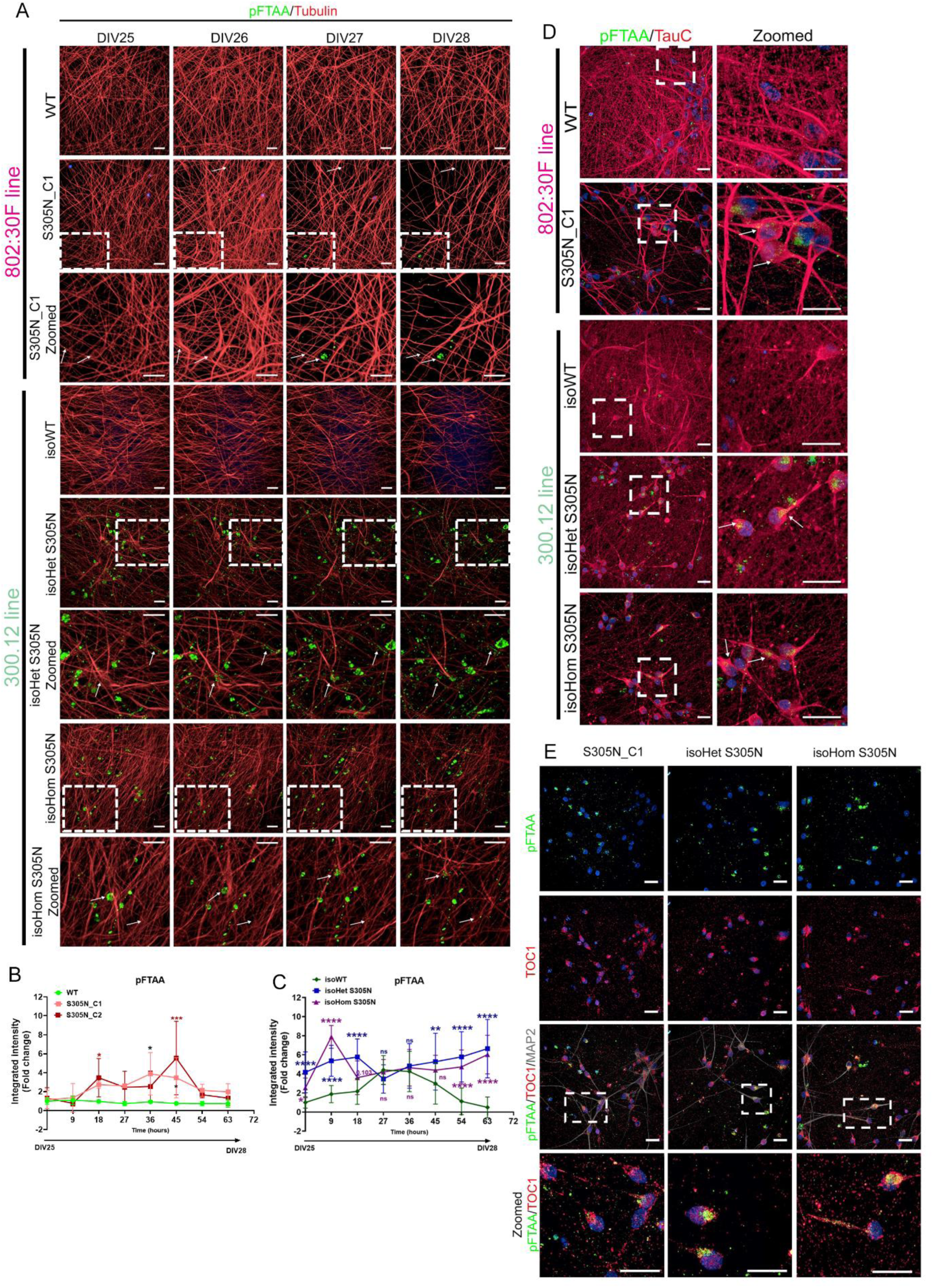
pFTAA positive inclusions in S305N i^3^N neurons. (A) Representative live-cell images of i3N neurons labelled with pFTAA and Tubulin over time. Dyes were added at DIV25, and images were acquired every 9 hours. (B) Quantification of pFTAA signal within Tubulin-positive regions (surrounding the nuclei) over time in WT and S305N_C1 and C2 neurons (two-way ANOVA followed by Tukey post-hoc test; mean ± SD, n = 3 independent neuronal differentiations). (C) Quantification of pFTAA signal within Tubulin-positive regions over time in isoWT, isoHet, and isoHom S305N neurons (two-way ANOVA followed by Tukey post-hoc test; mean ± SD, n = 3 independent neuronal differentiations). (D) Representative images of DIV28 neurons fixed with methanol and co-labelled with pFTAA and TauC. (E) Representative images showing colocalization of pFTAA with TOC1 and MAP2 in DIV28 neurons fixed with methanol. Statistical significance: *p < 0.05; **p < 0.01; ***p < 0.001; ****p < 0.0001.

As pFTAA is not specific for tau, we confirmed that the pFTAA signal seen in the S305N cells represented tau assemblies by co-staining methanol-fixed neurons with TauC (total tau) and TOC1 (oligomeric tau). In all S305N-mutant lines, pFTAA signal colocalised with TauC, and it was predominantly perinuclear (Fig. 6D and Fig. S8A for S305N_C2). As both TOC1 and pFTAA can label dying neurons, we co-stained with MAP2 to assess neuronal viability. Most pFTAA- and TOC1-positive neurons also labelled positively for MAP2, suggesting they were still viable and part of the neuronal cell population (Fig. 6E and Fig. S8A for S305N_C2). Further, colocalization of pFTAA with Lysotracker Red in S305N_C1, S305N_C2, isoHet and isoHom S305N neurons revealed that tau assemblies were accumulating in lysosomes at DIV28, rather than being degraded (Fig. S8B). These results demonstrate that tau assemblies in S305N neurons develop progressively over time, with isoHet and isoHom neurons exhibiting earlier and more extensive aggregate formation than S305N_C1 and C2. The pFTAA-positive structures likely represent early oligomeric and potentially protofibrillar tau species that accumulate in tubulin-rich compartments and evade lysosomal degradation, contributing to tau pathogenesis in S305N mutant neurons.

### Cytoskeletal alterations in S305N neurons

Given the known role of tau in microtubule dynamics, we assessed post-translational modifications (PTMs) of tubulin using immunocytochemistry and immunoblotting. Tyrosinated α-tubulin (Tyr-Tubulin), a marker of dynamic and newly polymerized microtubules (*47, 48*), was more prominently localized in the somatodendritic compartment of S305N-mutant neurons (Fig. 7A). Quantitative analysis revealed a significant increase in Tyr-Tubulin signal within MAP2-positive regions of S305N_C1 and C2 neurons at DIV21, whereas at DIV28 only S305N_C1 presented a close to significant change (p=0.06), with isoHom neurons showing elevated levels only at DIV28 (Fig. 7B, C). Longitudinal quantification of Tyr-Tubulin levels showed a progressive decline in WT and isoWT neurons, consistent with the transition toward stable axonal architecture during maturation. In contrast, S305N lines with a homozygous mutation exhibited elevated Tyr-Tub levels compared to their respective WT controls, suggesting altered microtubule dynamics or delayed axonal stabilization (Fig. 7D-F). We also investigated tubulin polyglutamylation (PolyE), a PTM that enhances microtubule rigidity and resistance to depolymerization (*49*). In WT and isoWT neurons, PolyE Tubulin levels increased over time, consistent with the formation of stable microtubules. However, S305N_C1 and C2 neurons start presenting a decrease in PolyE Tubulin from DIV17; this change was only significant at DIV28. The isoHet and isoHom S305N neurons had significantly lower levels of polyE Tubulin from DIV24, and a change was only statistically significant in the isoHet line (Fig. 7D-H). Although the cytoskeleton changes were only observed in the mutant lines, the precise relationship with tau type and distribution will require more extensive testing.

**Fig. 7.**
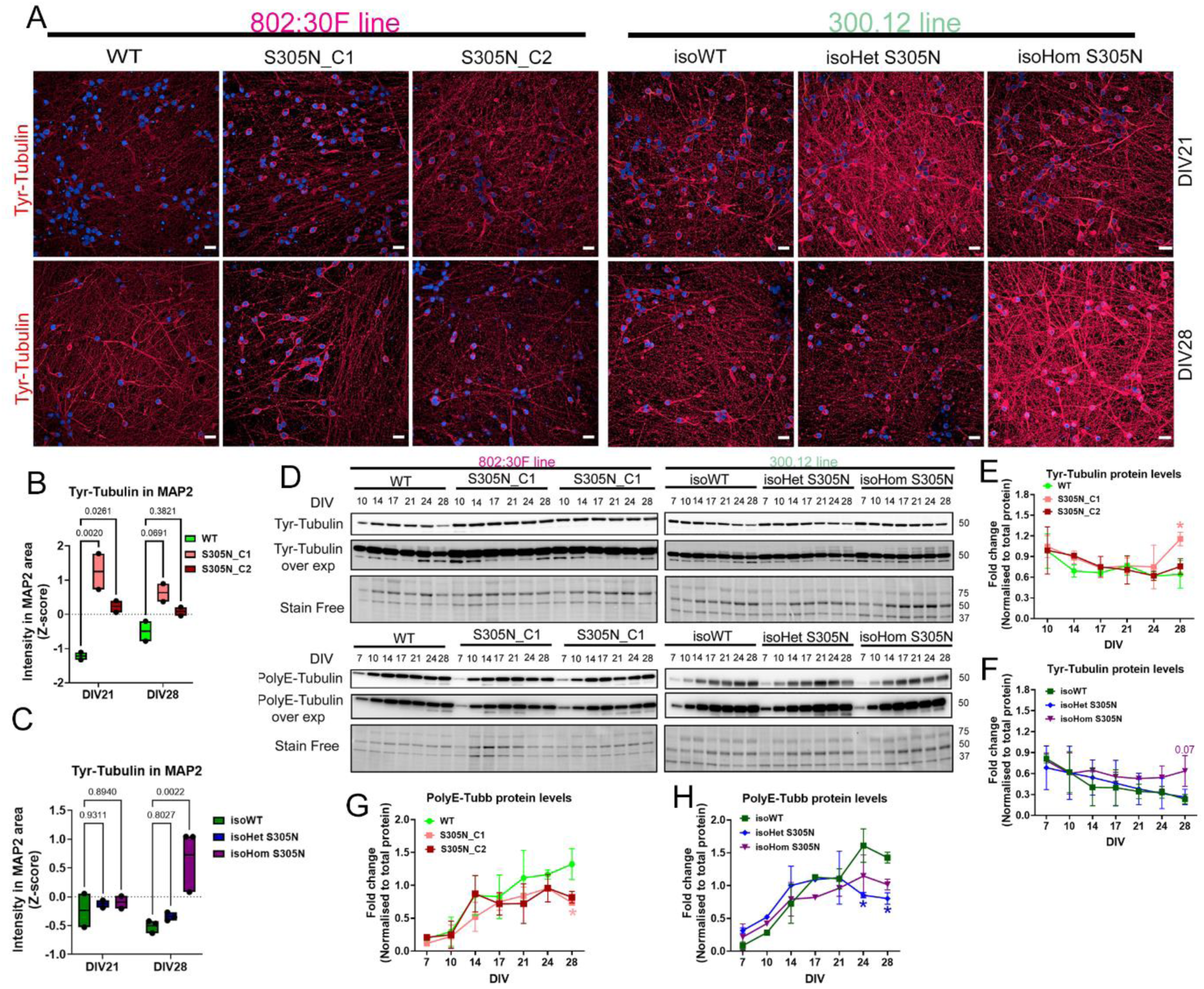
S305N i^3^N neurons demonstrate markers of cytoskeleton change. (A) Representative immunostaining of tyrosinated α-tubulin (Tyr-Tubulin), in i^3^N neurons at DIV21 and DIV28 across all neuronal lines. (B) Quantification of Tyr-Tubulin signal within MAP2-positive regions in WT versus S305N_C1 and S305N_C2 neurons at DIV21 and DIV28. Statistical analysis was performed using two-way ANOVA followed by Tukey’s post-hoc test (mean ± SD, n = 2 independent neuronal differentiations). (C) Quantification of Tyr-Tubulin signal within MAP2-positive regions in isoWT versus isoHet and isoHom S305N neurons at DIV21 and DIV28. Statistical analysis was performed using two-way ANOVA followed by Tukey’s post-hoc test (mean ± SD, n = 2–3 independent neuronal differentiations). (D) Representative immunoblots showing Tyr-Tubulin (top panel) and polyglutamylated tubulin (PolyE Tubulin; bottom panel) expression over time in all neuronal lines. (E) Quantification of Tyr-Tubulin protein levels in WT versus S305N_C1 and C2 neurons over time, normalized to total protein. Statistical analysis: two-way ANOVA with Tukey’s post-hoc test (mean ± SD, n = 2). (F) Quantification of Tyr-Tubulin protein levels in isoWT versus isoHet and isoHom S305N neurons over time, normalized to total protein. Statistical analysis: two-way ANOVA with Tukey’s post-hoc test (mean ± SD, n = 2). (G) Quantification of PolyE Tubulin levels normalized to total protein over time in (G) WT versus S305N_C1 and C2 neurons and (H) in isoWT versus isoHet and isoHom S305N neurons. Statistical analysis: two-way ANOVA with Tukey’s post-hoc test (mean ± SD, n = 2). Statistical significance: *p < 0.05.

### A novel neuronal model that responds to tau modulators

A major limitation in the field is the lack of a high-throughput screening assay to monitor the levels of endogenous tau in human iPSC-derived neurons. To address this, we inserted an 11–amino acid peptide (HiBiT) that allows sensitive luminescence-based detection of tagged proteins when complemented with LgBiT in a lytic luminescence assay. CRISPR/Cas9-mediated genome editing was used to insert the HiBiT tag at the C-terminus of the MAPT gene in S305N_C1, which was chosen as it presented the strongest phenotype regarding seed-competency and TOC1/pFTAA accumulation. Successful integration and expression of HiBiT-tagged tau was confirmed by immunocytochemistry in methanol-fixed neurons using Tau13 and TauC antibodies, both of which colocalised with the HiBiT signal at DIV21 (Fig. 8A). To test the utility of the assay, we treated neurons with a panel of compounds known to affect tau dynamics, including the autophagy inhibitor bafilomycin A1 (Baf) the mTOR inhibitor KU-0063794 (KU), and the small molecule Anle138b, reported to inhibit tau aggregation (*50*). After 3 hours of treatment, Baf led to a 50% increase in HiBiT signal, while KU led to a reduction. These effects were more pronounced after 20 hours, with Baf significantly increasing HiBiT signal by almost 2-fold and both KU and Anle138b showing more than 30% reduction (Fig. 8B) compared to DMSO-treated cells. Based on this, we selected the 20-hour timepoint for subsequent screening. The HiBiT lytic assay only monitors the amount of HiBiT-tagged tau, and not the abundance or cellular distribution of tau proteoforms, which are important aspects of tau pathogenesis. Characterisation of S305N_C1 i^3^N neurons had revealed that tau relocated from axonal to somatodendritic compartments (Fig. 2B), a redistribution that is known to increase its propensity for aggregation due to reduced microtubule binding. Compounds that can prevent this or modulate the levels of tau overall may be therapeutically relevant. We therefore designed a dual readout platform combining the HiBiT luminescence assay with high-content imaging for phenotypic screening. This setup allows parallel assessment of compound effects on tau levels (via the lytic HiBiT assay), tau localisation (via TauC immunostaining) and cell viability (Fig. 8C). To validate the platform, we treated S305N_C1 neurons with Baf or KU for 20 hours. TauC staining confirmed increased tau accumulation with Baf, including a potential increase in axonal tau, while KU treatment resulted in reduced tau staining, consistent with enhanced clearance (Fig. 8D). Dose-response experiments with KU and Baf showed concentration-dependent modulation of the HiBiT signal (Fig. 8E and F), demonstrating the assay’s sensitivity and dynamic range. Notably, neither compound adversely affected cell viability under these conditions; in fact, high concentrations of KU slightly increased viability (∼25%), suggesting a potential protective effect. Collectively, these results demonstrate that the HiBiT-tagged S305N_C1 neuronal model provides a robust platform for quantifying changes in tau levels and cellular distribution, for the screening of tau-modulating compounds in a high-throughput manner.

**Fig. 8.**
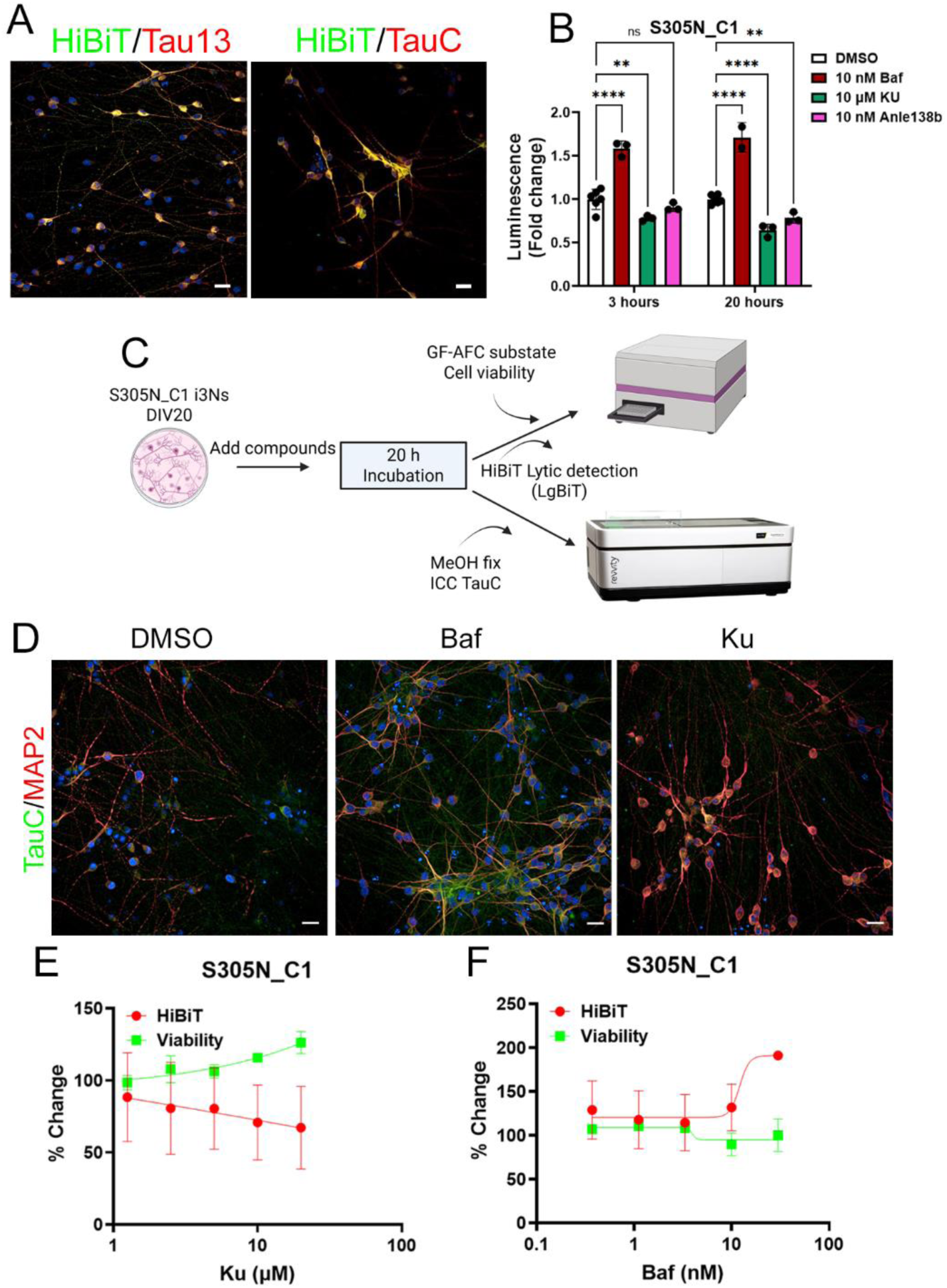
Optimisation of a drug screen assay for tau modulators. (A) Representative images showing co-localization of the HiBiT tag with Tau13 (left) and TauC (right) in DIV21 neurons. (B) Luminescence changes following treatment with DMSO, 10 nM bafilomycin A1 (Baf), 10 μM KU-0063794 (Ku), or 10 nM Anle138b for either 3 or 20 hours at DIV21. Statistical analysis was performed using two-way ANOVA followed by Tukey’s post-hoc test (mean ± SD, n = 2 independent neuronal differentiations). (C) Schematic representation of the proposed screening platform for 4R tau modulators (created with BioRender). (D) Representative images of methanol-fixed DIV21 neurons stained with TauC and MAP2 after 20-hour treatment with DMSO, 10 nM Baf, or 10 μM Ku. (E–F) Dose-response curves showing changes in luminescence (HiBiT signal) and cell viability in DIV21 S305N_C1 neurons treated with Ku (E) or Baf (F) for 20 hours. Data represent mean ± SD from 2 independent neuronal differentiation experiments, with three wells per condition per experiment. Statistical significance: **p < 0.0, ****p < 0.0001.

## Discussion

The human neuron cell lines we describe here represent a significant advance for the neurodegeneration field as they recapitulate several of the key pathogenic events observed in human diseases associated with tauopathy. One of the major problems in the field has been that most iPSC-derived human neurons, even those expressing FTD-causing mutations, model the fetal state and do not accumulate appreciable levels of 4R tau, and they do not form endogenous seed-competent tau assemblies. Human neuronal lines that do accumulate 4R tau required multiple tau mutations to increase 4R tau expression, or extended culturing time, and tau aggregates had to be induced by incubation with exogenous tau seeds, all of which limit their physiological relevance (*17–21*). Our data, from two independent human donor lines, demonstrate that expressing the S305N mutation leads to rapid, elevated 4R tau accumulation, which is associated with altered tau phosphorylation, distribution, aggregation, and downstream cellular pathways critical for neuronal homeostasis and microtubule stability (see Summary Table S1), recapitulating what is observed in a wide range of human tauopathies.

Clinically, S305N-linked FTLD-MAPT is associated with 4R tau pathology, ring-shaped perinuclear aggregated tau, and neurofibrillary tangles (*32, 40, 41*). Our neuronal lines recapitulate much of this phenotype (including seed-competent tau) and we believe they represent an early stage of this pathogenic process. Whether the effects observed in S305N mutant lines mirror the pathological events underlying other MAPT mutations, sporadic primary tauopathies, secondary tauopathies, and inflammation-driven tauopathies (*51*), or other neurodegenerative disorders remains unknown. However, this may be irrelevant if heterogeneous etiologies converge on the common pathological pathways of tau hyperphosphorylation and aggregation.

The levels of phosphorylated tau were significantly altered in the mutant neuronal lines, particularly in S305N neurons with the homozygous mutation. This was demonstrated by increased CP13 and/or AT8 immunoreactivity, regardless of overall reduced total tau levels. However, we observed that despite the overall relatively lower levels of 4R tau, isoHet S305N neurons presented a trend towards higher TOC1, pFTAA, and seed-competency compared to isoHom S305N, suggesting that the relationship between 4R levels, phosphorylation status and aggregate formation is not direct. Increased phosphorylation in the same isoHom line has been reported; however, the authors did not measure changes in seeding activity or accumulation of TOC1/pFTAA (*29*). We observed interesting differences in the distribution of several phosphorylated tau epitopes, with AT100 only being detected in the axons and not in the somatodendritic compartment. The AT100 (pThr212/pSer214) phosphorylation sites are thought to be critical for modulating the ability of tau to bind and maintain microtubule stability (*52–56*). Tau hyperphosphorylation at the AT100 epitope may promote, or indicate tau detachment from microtubules, increasing its propensity to form small soluble oligomers or insoluble aggregates, which could be toxic to neurons (*52–56*). AT100 was increased at DIV28 in all neuronal lines harbouring the S305N mutation, independent of the amount of total tau, and at a timepoint when pFTAA positive signal was present and neurons had formed endogenous, seed-competent tau. The presence of increased AT100 immunoreactive tau in axonal compartments might indicate that tau oligomer formation is initiated there, and that tau seeds are then transported to the soma, where we see the highest accumulation of TOC1 immunoreactive tau and pFTAA signal. AT8 is the only phosphorylated tau antibody located in the soma, and it is similar to TOC1/pFTAA distribution. Notably, longitudinal imaging revealed that tau assemblies accumulate in axons *prior* to the soma, suggesting that the axons are the initial point of vulnerability. The appearance of TOC1-positive oligomeric tau, particularly in the perinuclear region and in MAP2-positive compartments, indicates the presence of aggregation-prone tau species in these compartments. Similar structures are present in neurons in the cortex of human S305N brain tissue. The high density of i^3^N cultures limited our ability to consistently resolve pFTAA-positive puncta along axons; however, we observed discrete puncta predominantly near nuclei and within pre-axonal segments. At DIV28, pFTAA signal was largely confined to MAP2-positive regions and co-localized with TauC, confirming that pFTAA was labelling abnormal conformers of tau. The colocalization of pFTAA-positive structures with lysosomes suggests that these assemblies are resistant to degradation, consistent with the behaviour of oligomeric tau (*57, 58*). However, the absence of tau labelling by MC1, a conformation-specific antibody recognizing tau in neurofibrillary tangles, suggests that despite the accumulation of oligomeric, seed-competent tau in our models, the formation of more mature tangle-like pathology may require longer culture times, the presence of mixed neural and glial populations or additional stressors that more closely mimic the *in vivo* disease environment. The presence of tau seeds in DIV28 neuronal lysates, as shown by the biosensor assay, suggests that S305N neurons not only accumulate misfolded tau, but also generate seed-competent species capable of propagating pathology. The absence of seeding at DIV21, despite a detectable TOC1 signal, implies a temporal progression from early oligomeric intermediates to seed-competent assemblies. However, overall, the seeding signal was low, possibly due to quite low levels of tau in the neurons, which makes it difficult to identify small changes at DIV21 compared to DIV28. Of note, our longitudinal studies support a model where tau misfolding and phosphorylation precede the formation of seed-competent tau species.

Biochemical and morphological data, which include the presence of tubulin blebbing, reduced MAP2 levels, and reduced VGLUT1 expression, highlight the functional consequences of tau dysregulation in the i3N neurons. Post-translational modifications of tubulin provide additional insight into the cellular impact. Elevated tyrosinated tubulin and reduced polyglutamylation in the neurons with more than 75% 4R tau suggest a shift toward a more dynamic, less stable microtubule network. These changes are consistent with a failure to mature structurally stable axons. In human AD brain tissue, reductions in acetylated (*59*) and polyglutamylated (*49*) α-tubulin has been reported, suggesting that tau dysregulation affects microtubule structure both *in vitro* and *in vivo* (*60*).

One of the key current limitations for drug discovery targeting tauopathies is the absence of a high-throughput assay to quantify 4R tau dynamics and functional outcomes in living, human neurons. To address this, we developed a novel luminescence-based assay by knocking in a HiBiT tag at the endogenous *MAPT* C-terminus in the S305N_C1 iPSC line. This system enables rapid, sensitive detection of endogenous tau levels and offers the added benefit of compatibility with drug screening pipelines. Using the lytic assay, we validated the effects of known tau-modulating compounds. Bafilomycin A1 treatment increased HiBiT signal, reflecting tau accumulation, whereas KU-0063794 and Anle138b reduced tau levels in a time- and concentration-dependent manner. This system also allows phenotypic readouts through high-content imaging, enabling the identification of compounds that not only alter overall tau (C-terminus) levels, but also compounds that affect the subcellular localisation of different tau proteoforms, a key pathological feature of human tauopathies that is represented in our cell model. Notably, these lines are the first single coding mutation i^3^N neuronal line to report formation of endogenous tau assemblies, an important target of therapeutics, which can be monitored longitudinally in these lines. Based on our results, expansion of the toolbelt of S305N mutant lines (including the development of a KOLF line for cross-lab comparison), is now justified.

In conclusion, our study provides insight into how the accumulation of 4R human tau caused by the S305N mutation disrupts the normal biological function of tau, leading to pathological changes including altered tau, increased phosphorylation and aggregation, impaired microtubule stability, and dysregulation of key neuronal pathways. The reasonable timeframe and progressive nature of the tau pathology observed in these neuronal cell lines will make them a valuable tool for assessing the sequence of pathological events during the initial stages and early progression of 4R tauopathies. Additionally, being able to assess the order of events and the spatio-temporal relationship between cellular location and tau proteoform distribution, in the absence of overexpression artifacts or prolonged culturing, is a significant strength of this model. Our findings not only underscore the multifaceted role of tau in neuronal development and maintenance but also highlight potential therapeutic targets for the earliest stages of tau pathogenesis, and they provide proof-of-concept data for an assay platform to screen them.

## Limitations of the study

One limitation of our model is that we used the piggyBac system to introduce hNGN2 into the iPSC lines, which typically results in random integration into the genome. No karyotype abnormalities were observed, but there is still the possibility that hNGN2 could have disrupted some genes in these cells. Replicating some of these findings in either directly differentiated neurons or in iPSC lines where hNGN2 has been introduced into a safe-harbour location will further strengthen our understanding of how the S305N mutation affects neuronal differentiation and tau aggregation.

Two observations could impact the interpretation of the study. Firstly, the S305N_C1 and C2 neurons express 4R tau from a relatively early stage of neuronal differentiation, which could impact developmentally sensitive events. It is not known if 4R tau predominates during early life in humans expressing the S305N mutation. Secondly, there was a marked reduction in total tau protein levels in mutant lines compared to their isogenic WT controls. We do not believe that this is an artifact related to the donor line as the effect was seen in two clones from one of the lines, and in two different donor lines, with the isoHom S305N presenting a similar phenotype to S305N_C1 and C2 neurons, albeit at a slightly later timepoint (after DIV21). As this corresponds to the time when 4R tau levels increase by more than 50%, the shift toward excessive 4R tau is likely implicated in disrupted protein turnover and axonal destabilization. A previous study has shown that 4R tau isoforms have a quicker turnover rate than 3R tau in iPSC-derived cells (*61*), which could result in lower overall tau levels in the mutant lines compared to the predominently 3R-expressing WT line. Our observations are consistent with those published previously for the iso lines (*29*).

## Resource availability

The data generated and/or analysed in this study is available from the authors upon request.

Cell lines 802:30F S305N_C1 and C2 are new lines generated in this study and are available upon request.

## Acknowledgments

This work was mainly supported by the Rainwater Charitable Foundation as well as by the UK Dementia Research Institute through UK DRI Ltd, principally funded by the Medical Research Council. We would like to thank NCRAD for the patient-derived S305N lines (300.12) and Dr. Kathryn Bowles (UK DRI Edinburgh) for generating isogenic lines and her advice on using the cells. We thank Prof. Selina Wray (UCL) and her team for their help and advice on setting up the iPSC cultures, the ARUK Drug Discovery Institute (UCL) for access to equipment and resources, the ALBORADA Drug Discovery Institute (Cambridge) for advice on developing the drug screening assay, Dr. Cristina D’Abramo (Peter Davies lab, the Feinstein Institutes for Medical Research, New York) for CP13 and MC1 antibodies, Dr. Nicholas Kanaan (Lester Binder lab Northwestern University) for TOC1 antibody, Dr. Brad Boeve and The Mayo Clinic for human S305N tissue, Dr. David Villaroel (Dr. James Sleigh lab) for the tubulin antibodies and advice and Dr. Michael Ward (NIA) for the hNGN2 piggyBac plasmids. Dr. Malcolm Macleod (Edinburgh University) reviewed this work from the perspective of data, image and methodological integrity.

## Author contributions

K.E.D conceived the project and E.T designed and performed the experiments. S.B, T.B. M.F, N.W, S.C helped perform experiments and E.T M.B, D.G, E.Tu analysed data. R.C and A.I provided protocols and plasmids. E.T and K.E.D wrote the manuscript. All authors discussed, reviewed, and edited the manuscript.

## Key resources table

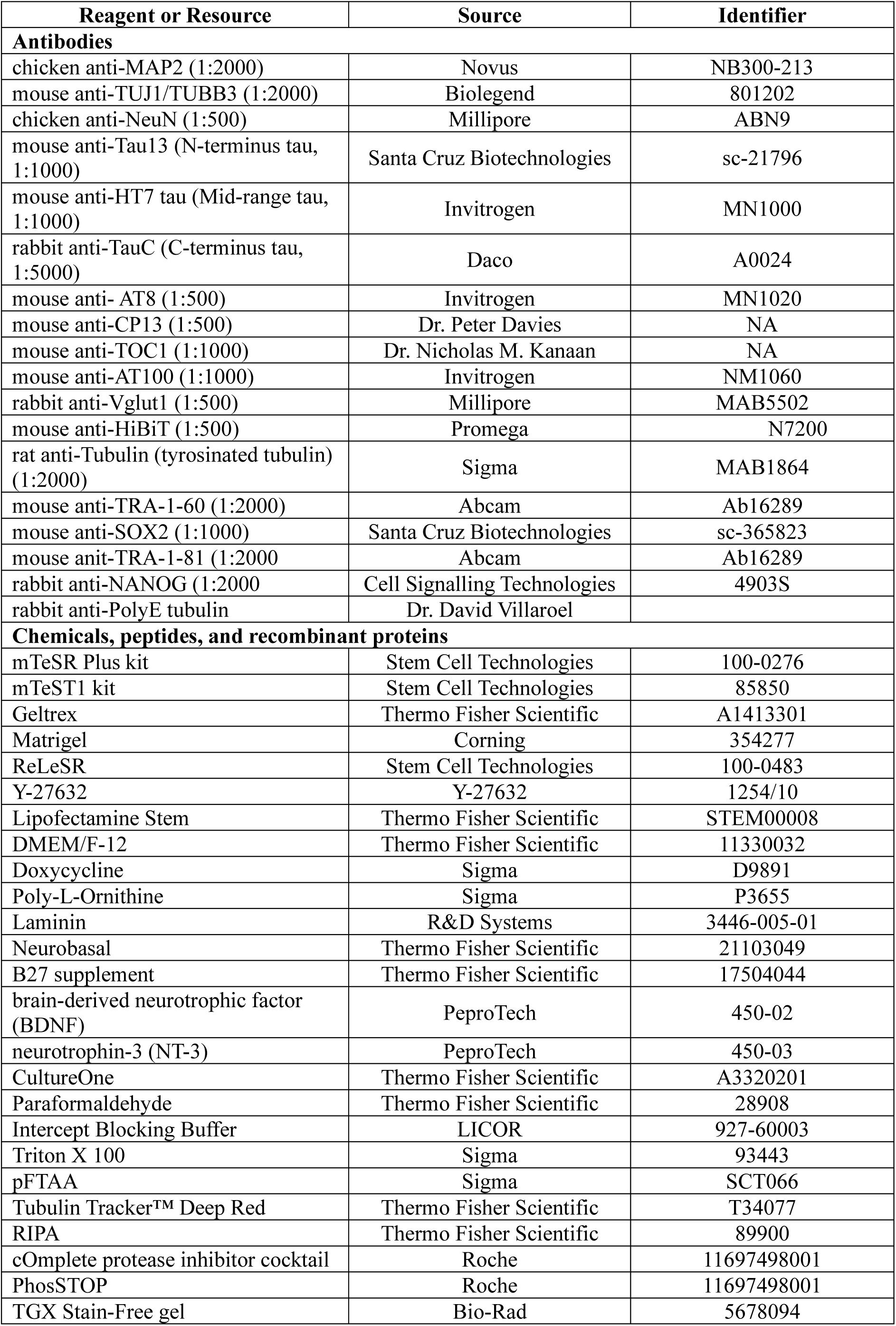
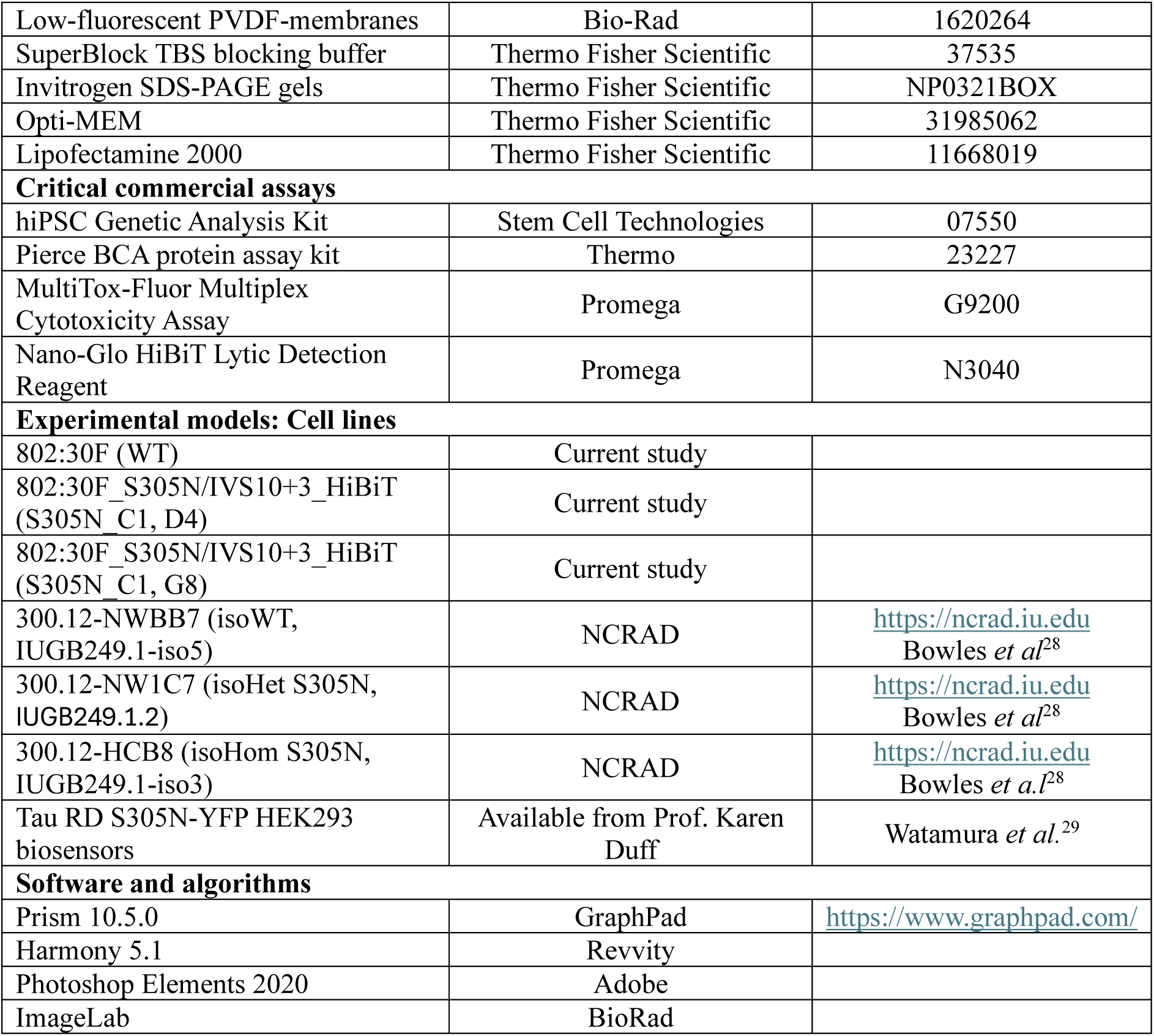

## Methods

### Generation of iPSC line with a S305N/IVS10+3 mutation via CRISPR-Cas9

iPSCs with mutations S305N and IVS10+3 were generated by Synthego utilising one of their in-house iPSC lines (802:30F). Briefly, a single guide (GUACUCACACUGCCGCCUCC) was utilised to introduce both mutations by using their in-house CRISP-Cas9 editing protocol. Sanger sequencing was utilised to identify clones with homozygote double mutation. Consequently, one of the clones (Clone B11) carrying the double homozygote mutation was selected to knock in a HiBiT tag in the C-terminus of *MAPT*. Two clones were provided (D4 and G8) by Synthego that contained the double mutant and the HiBiT tag (S305N_C1 and C2) as well as the mock-transfected wild-type cells (WT). The top 3 off-target sites for both single guide and HiBiT tag were assessed via PCR and sequencing. No off-target sites were identified.

### Culture of iPSCs

iPSCs on the 802:30F background were maintained in Geltrex (Thermo; A1413301) coated plates using mTeSR Plus (StemCell Technologies; 100-0276) and passaged using 0.5 mM EDTA (Thermo; 15575-020). Patient-derived iPSCs on the 300.12 background were obtained from NCRAD. These cells were maintained in Matrigel (Corning, 354277) using mTeSR1 (StemCell Technologies; 85850) and passaged using ReLeSR (Stem Cell Technologies; 100-0483). All iPSC lines for this work were regularly tested for mycoplasma.

### Stable integration of piggyBac plasmids into iPSCs

To generate stably expressing hNGN2-expressing iPSC we integrated a BFP-containing doxycycline-induced hNGN2 cassette using the piggyBac system. iPSCs were washed with PBS, dissociated with Accutase (Gibco; A1110501), and plated into coated plates (Geltrex or Matrigel) at 1x10^6^ cells/well of a 6-well plate in mTeSR Plus or mTeSR1 containing 10 μM Y-27632 (Tocris; 1254/10). Once cells were attached, they were washed with PBS before being transfected with constructs containing the donor doxycycline-induced hNGN2 and BFP fluorescent reporter flanked by transposon terminal repeats and the piggyBac transposase. For the transfection, 3 μg of total plasmid DNA was used (2:1 donor to transposase ratio), with Lipofectamine Stem (Thermo; STEM00008) as per the manufacturer’s instructions. Forty-eight hours post-transfection, media was changed and supplemented with 1 μg/mL puromycin for 3 days to select for cells that had successfully integrated the hNGN2. To obtain a pure population of cells with high levels of BFP expression due to cassette insertion, cells were subjected to fluorescence-activated cell sorting (FACS) using BFP fluorescence. To ensure genomic stability, iPSCs were karyotyped using low-coverage sequencing (UCL Genomics) with 50 kbp resolution. All lines were also checked routinely for abnormalities using the hiPSC Genetic Analysis Kit (Stem Cell Technologies, 07550).

### iPSC differentiation into cortical i^3^N neurons

Induction of iPSC into cortical neurons was initiated by dissociating cells in Accutase and plating 1x10^6^ iPSC onto Geltrex (Synthego 802:30F lines) or Matrigel (NCRAD 300.12 lines) coated 6-well plates using DMEM/F-12 with Glutamax/Hepes (Thermo, 11330032) containing 2 μg/mL doxycycline (Sigma; D9891), 1X N2 Supplement (Thermo, 17502048), 1X non-essential amino acids (Thermo, 11140050) and 10 μM Y-27632. The media was replaced daily for the next 2 days with media without Y-27632. At day 3, induced cells were dissociated with Accutase and replated into plates double-coated with 0.1 mg/mL Poly-L-Ornithine (PLO; Sigma, P3655; 24 hours at 37°C) and 30 μg/mL Laminin (R&D Systems, 3446-005-01; 24-72 hours at 37°C). Cells were plated into 6-well plates at 0.6x10^6^ cells/well, 24-well plates at 0.125x10^6^ cells/well, into 96-well Phenoplates (Revvity; 6055300) or white plates for Luminescence assays (Greiner, 655083) at 30,000 cells/well. Neuronal maturation media consisted of Neurobasal (Thermo, 21103049) media containing 1X B27 (Thermo, 17504044), 10 ng/mL brain-derived neurotrophic factor (BDNF, PeproTech, 450-02), 10 ng/mL neurotrophin-3 (NT-3, PeproTech, 450-03), 6-12 μg/mL Laminin, 2 μg/mLl doxycycline, and 1X CultureOne (Thermo, A3320201). Once a week, one-half of the media was replaced with fresh media until cells were collected.

### Immunocytochemistry, imaging, and analysis

Human iPSCs were fixed with 4% methanol-free paraformaldehyde (Thermo; 28908) diluted in PBS for 15 min at room temperature and washed 3 times in DPBS (Thermo; 14190-094). Most of the i^3^N neurons were fixed with 100% methanol for 15 min at -20° C. Cells were washed twice with ice-cold DPBS. Cells were blocked/permeabilised with Intercept Blocking Buffer (LICOR; 927-60003) containing 0.1% Triton (Sigma; 93443) for 1 hour at room temperature. Primary antibodies diluted in blocking buffer were added to the cells and incubated overnight at 4°C and then washed twice with DPBS. The following primary antibodies were used: chicken anti-MAP2 (1:2000, Novus; NB300-213), mouse anti-TUJ1/TUBB3 (1:2000, Biolegend; 801202), chicken anti-NeuN (1:500, Millipore; ABN9), mouse anti-Tau13 (N-terminus tau, 1:1000, Santa Cruz Biotechnologies; sc-21796), mouse anti-HT7 tau (Mid-range tau, 1:1000, Invitrogen; MN1000), rabbit anti-TauC (C-terminus tau, 1:5000, Daco; A0024), mouse anti-AT8 (1:500, Invitrogen; MN1020), mouse anti-CP13 (1:500, Peter Davies), mouse anti-TOC1 (1:1000, Nicholas M. Kanaan) mouse anti-AT100 (1:1000, Invitrogen; NM1060), rabbit anti-VGLUT1 (1:500, Millipore; MAB5502), mouse anti-HiBiT (1:500, Promega, N7200), rat anti-Tubulin (tyrosinated α-tubulin) (1:2000, Sigma; MAB1864), mouse anti-TRA-1-60 (1:2000, Abcam; Ab16289), mouse anti-SOX2 (1:1000, Santa Cruz Biotechnologies; sc-365823), mouse anit-TRA-1-81 (1:2000, Abcam; Ab16289) and rabbit anti-NANOG (1:2000, Cell Signalling Technologies; 4903S). Secondary antibodies diluted in blocking buffer, including goat anti-mouse, anti-rabbit or anti-chicken conjugated with Alexa Fluor 488, Alexa Fluor 568 or Alexa Fluor 647 (1:1000) as well as Hoechst (1:1000, Thermo; 33342) were added into the cells and incubated for 1 hour at room temperature. Cells were washed twice with DPBS.

Images were acquired with the high-content microscope Opera Phenix Plus (Revvity) by using the 40x water immersion objective with 12 fields of view per well and z-stack of 6 planes (0.5 μm per plane). Acquisition settings (exposure times and laser power) were adjusted to be above the camera background and kept below saturation levels. Imaging settings were kept the same between biological replicates.

Acquired images were analysed in Harmony 5.1. Briefly, a stack of each image was maximum projected and basic flatfield correction was applied. Cells were segmented by identifying the nuclei of live cells only (‘find nuclei building block’). ‘Find image region’ was used to identify tau or Tubulin in the imaged field. For antibodies where staining was localised in the soma, such as TOC1 or AT8, MAP2 was used to identify cytoplasm and consequently ‘find image region with absolute thresholding’ was used to identify the TOC1 positive area. In some cases, TOC1 labelled dead cells, and a filter was applied to remove these areas during the analysis. Both intensity and area of the studied protein were calculated, and the results were normalised to the number of cells. Integrated intensity was calculated by multiplying the area by the mean intensity. Due to variation in the absolute values of the integrated intensity between biological replicates, Z-scoring was used instead. For each biological experiment, the Z-score was calculated with the formula 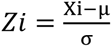, where Xi is the value of the individual technical replicate, μ is the mean of all conditions for the specific biological experiment and σ is the standard deviation for all conditions for the specific biological experiment. Average Z-scores from different biological replicates were plotted in GraphPad Prism.

### pFTAA live imaging of i^3^N neurons and imaging analysis

To monitor pFTAA staining in i^3^N neurons, cells were grown until DIV25. Old media was aspirated, and cells were washed with fresh Neurobasal media without Phenol Red (Thermo; 12348017) containing 1X B27 (Thermo, 17504044), 10 ng/mL brain-derived neurotrophic factor (BDNF, PeproTech, 450-02), 10 ng/mL neurotrophin-3 (NT-3, PeproTech, 450-03), 6-12 μg/mL Laminin, 2 μg/mL doxycycline and 1X CultureOne (Thermo, A3320201) to remove dead cell from the culture. Fresh neuronal media containing 1.5 μM pFTAA (Sigma, SCT066) and 1:10000 Tubulin Tracker™ Deep Red (Thermo, T34077) was added and plates were incubated for 30 min before starting live imaging on the high-content microscope Opera Phenix Plus by using the 63x water immersion objective with 9 fields of view per well and z-stack of 6 planes (0.5 μm) with the environmental control enabled (5% CO_2_, 37° C and humidity). Images were acquired every 9 h for four continuous days. The pFTAA signal was detected by using 405nm/550nm Ex/Em filters and Tubulin at 647nm/665nm Ex/Em, BFP was also imaged as a nuclei marker (375nm/450nm Ex/Em, it is part of the hNGN2 plasmid). For experiments where we fixed cells after labelling with pFTAA, a similar approach was followed; however, plates were maintained in the incubator before fixation with methanol and immunostaining.

Acquired images were analysed in Harmony 5.1. Briefly, images were maximum projected, and basic flatfield correction was applied. Live cells were identified by the ‘find nuclei’ (using the BFP channel) function, bead cells were filtered out using a different approach such as nuclei size and intensity. The cytoplasm of living cells was then identified using the tubulin channel. ‘Find regions building block’ was used to identify pFTAA positive areas by using the absolute threshold option to make sure that we detected objects above background (600-700 was used as the lowest threshold). Objects where pFTAA was colocalised with dead cells were excluded from the analysis by filtering them out. Both pFTAA area and intensity were calculated and normalised to the number of live cells. Results are presented as integrated intensity, and data were plotted on GraphPad Prism.

### Quantitative PCR (qPCR)

iPSC samples for RNA extraction were collected by washing cells with PBS and dissociating cells with either EDTA or ReLeSa, spinning down at 300xg for 7 min at 4°C. Pellets were stored at -80°C. i^3^Ns were washed with PBS before adding fresh PBS and lifting the monolayer by swirling the plate. Cells were collected and spun down at 300xg for 7 min at 4°C and pellets were stored at -70°C. RNA extraction was performed using ReliaPrep RNA Cell Miniprep System (Promega; Z6012) according to the manufacturer’s instructions. Reverse transcription was performed by using 300 ng RNA and LunaScript RT Supermix Kit (New England Biolabs; E3010) by following the manufacturer’s instructions. qPCR was performed using the Luna Universal qPCR Master Mix kit (New England Biolabs; M3003S), according to the manufacturer’s instructions and reactions were carried out using the Lightcycler 480 II (Roche). Expression values were calculated in Excel using the Ct method, where GAPDH was used as a control.

### Semi-quantitative PCR (sq-PCR)

Semi-quantitative PCR was conducted to quantify the ratio of 3R:4R mRNA. cDNA was used as a template, which amplified by primers (*23*) flanking exon 10 (forward 5^’^-AAGTCGCCGTCTTCCGCCAAG-3^’^; reverse 5^’^ - GTCCAGGGACCCAATCTTCGA-3^’^). Q5 2x master mix (New England Biolabs; M0492S) was used for the amplification, and PCR products were run on a 2% agarose gel with 381bp and 288bp fragments indicating 4R and 3R, respectively. Ratios were calculated as previously described (*62*), where ImageJ box plots and measure plots were used. Sum pixel intensity values were exported into Excel, and the percentage change was calculated by dividing each isoform value by the summed total intensity.

### Immunoblot analysis

All iPSC lines were differentiated into neurons on 6-well plates and samples were collected at DIV7, 10, 14, 17, 21, 24, 28 post-induction. Briefly, cells were washed with DPBS once before adding RIPA (Thermo; 89900) buffer containing cOmplete protease inhibitor cocktail (Roche; 11697498001) and PhosSTOP (Roche; 11697498001). Lysates were incubated on ice for 20 min, centrifuged at 12,000 g for 10 min at 4°C, and supernatants were collected. Samples for dephosphorylation assay were lysed in 50 mM Tris pH 7.6, 0.15 M NaCl, cOmplete protease inhibitor cocktail. Protein concentration was determined by using Pierce BCA protein assay kit (Thermo; 23227) for all lysates. Equal amounts were loaded into 4-20% TGX Stain-Free gels (Bio-Rad; 5678094) or dephosphorylated with lambda-phosphatase (Santa Cruz Biotechnology; sc-200312A). Proteins were transferred on to low-fluorescent PVDF-membranes (Bio-Rad; 1620264) blocked with 5% milk in TBS-Tween (0.02% Tween). Primary antibodies were diluted in SuperBlock TBS blocking buffer (Thermo Fisher Scientific; 37535) and incubated overnight at 4°C. Primary antibodies used included: mouse anti-Tau13 (N-terminus tau, 1:1000, Santa Cruz Biotechnologies; sc-21796), mouse anti-HT7 tau (Mid-range tau, 1:1000, Invitrogen; MN1000), rabbit anti-TauC (C-terminus tau, 1:5000, Daco, A0024), mouse anti-AT8 (1:500, Invitrogen, MN1020), mouse anti-CP13 (1:500, Peter Davies), rat anti-Tubulin (tyrosinated tubulin) (1:2000, Sigma; MAB1864), rabbit anti-PolyE tubulin (1:5000, gift from David Villaroel), anti-β-actin-FITC (Millipore, F3022). Incubation with AT8 antibody was not compatible with the TGX Stain-Free gel for that reason, 4-12% SDS-PAGE gels (Invitrogen, NP0321BOX) were used. Secondary HRP-conjugated goat anti-rabbit or goat anti-mouse antibodies were diluted in 5% milk in TBS-Tween for 1 hour at room temperature. Chemiluminescence (BioRad) was used for detection of immunoblotting and bands were quantified by intensity using ImageLab (BioRad). For the current study all blotted membranes were stripped by using PLUS Western Blot Stripping Buffer (Thermo Fisher, 10016433) for 12 min before incubating with a new primary antibody.

### Sample preparation for seeding with i^3^N neurons

i^3^N neurons were plates into 6-well plates and samples for seeding were collected at DIV21 and DIV28. A whole 6-well plate was used for these experiments. Cells were washed with DPBS before adding fresh DPBS and lifting cells as a monolayer by swirling the plate. Cells were pelleted by centrifuging at 300g for 7 min at 4°C and stored at -80°C. Pellets were lysed with 50 mM Tris pH 7.6, 0.15 M NaCl, cOmplete protease inhibitor cocktail. Samples were sonicated with a water bath sonicator (Qsonica Q800R3) at 4°C for 5 min at 65 amplitude (30 seconds on, 60 seconds off) before incubating on ice for 30 min. Samples were spun down at 300g for 5 min at 4°C and supernatant was collected. Protein concentration was determined by using Pierce BCA protein assay kit.

### Biosensor cell culture method and seeding

Tau RD S305N-YFP HEK293 biosensors and HEK293 Tau RD P301S FRET biosensor cells (ATCC CRL-3275) were cultured in DMEM (Thermo Fisher, cat. no. 41966-029) with 10% FBS and 1% Pen-Strep. Cells were plated at a density of 33,000 cells/well in a 96-well plate either PhenoPlates (Revvity) or standard tissue culture plates, in a volume of 130 μl medium per well. After 18 hours, cells were transduced with seeds; 1.25 μl Lipofectamine 2000 (Thermo Fisher Scientific, 11668019) was mixed with 8.75 μl Opti-MEM (Thermo Fisher Scientific, 31985062) and incubated at room temperature for 5 min before mixing with 18 μg of total protein of i^3^N total lysates. Complexes were incubated for 30 min at 37°C before transferring 20 μl to each well and further incubated with the cells for 72 hours. For each condition, three technical replicates were included.

### Imaging analysis for the S305N biosensor line

In S305N biosensors, 1:10000 Hoechst was added to the cells before imaging, and incubated for 10 min. Plates were imaged on the high-content microscope Opera Phenix Plus using the 20x water immersion objective with 20 fields of view per well and a z-stack of 8 planes (0.8 μm) with the environmental control enabled (5% CO_2_, 37°C, and humidity). Excitation wavelengths and emission filters were used as follows: YFP 488 nm/527-530 nm, Hoechst: 375 nm/435–480 nm. The acquired images were analysed in Harmony 5.2. Briefly, images were loaded as maximum projection and basic flatfield correction was applied. Cells were segmented by using the “find nuclei” building block using the Hoechst channel, consequently, “find cytoplasm” was identified by using the watershed of the Hoechst channel. The YFP positive tau inclusions were identified by using the “find spots” building block (method C) on the YFP channel. The analysis was exported, and the number of spots per nuclei was calculated.

### Flow Cytometry analysis for P301S biosensor line

To quantify tau seeding in the P301S FRET biosensor, cells were harvested using 0.25% trypsin, fixed in 4% paraformaldehyde for 10 minutes, and resuspended in flow cytometry buffer (1X PBS with 1 mM EDTA). FRET flow cytometry was performed using an LSR Fortessa Flow Cytometer (BD Biosciences). FRET-positive cells were identified as previously described(*43*). In brief, single cells double-positive for CFP and YFP were gated, and FRET-positive events were quantified within this population. The percentage of FRET-positive cells was calculated as output metric. Data were analysed using FCS Express v7 (De Novo Software) and GraphPad Prism.

### HiBiT Lytic detection assay and Viability assay

For the lytic assay, neurons were grown in white, Cellstar plates until DIV20 at 100 μl maintenance media. Neurons were treated at the indicated timepoint. To assess the neuronal viability at the final timepoint GF-AFC substrate from MultiTox-Fluor Multiplex Cytotoxicity Assay (Promega, G9200) was used following the manufacturer’s protocol. Briefly, 20 μl from the 5x GF-AFC substrate was added into the cells, incubated for 30 min at 37 °C before reading fluorescence (400Ex/505Em) at the FLUOstar Omega plate reader (BMG Labtech). Afterwards, 120 μl of the Nano-Glo HiBiT Lytic Detection Reagent (Promega, N3040) was added directly to the cells and incubated for 10 min before recording luminescence on a FLUOstar Omega plate reader with 0.2 s integration time.

### Human tissue

Post-mortem human brain tissue from an S305N carrier was kindly provided by The Mayo Clinic.

### Immunohistochemistry

For human FFPE staining, 8 μm sections were deparaffinized in xylene and rehydrated using graded alcohols. Following pressure cooker pretreatment in citrate buffer for 10 min and endogenous peroxidase quenching for 10 min, sections were blocked in 10% dried milk solution. Tissue sections were incubated with primary antibodies for 1 hour at room temperature, followed by biotinylated secondary incubation for 30 min at room temperature and avidin-biotin complex for 30 min. Colour was developed with 3,3′-diaminobenzidine/H_2_O_2_. Sections were scanned on a NanoZoomer Digital Pathology C9600 (Hamamatsu Photonics).

### Statistical analysis

All statistical analysis was conducted at GraphPad Prism. T-test was used to compare two groups and One-way ANOVA for three groups or more, with the appropriate post-hoc test as stated in the figure legend. Two-way ANOVA was used for comparison between cell lines and time with the appropriate post-hoc test.

## Supplementary Figures

**Fig S1.**
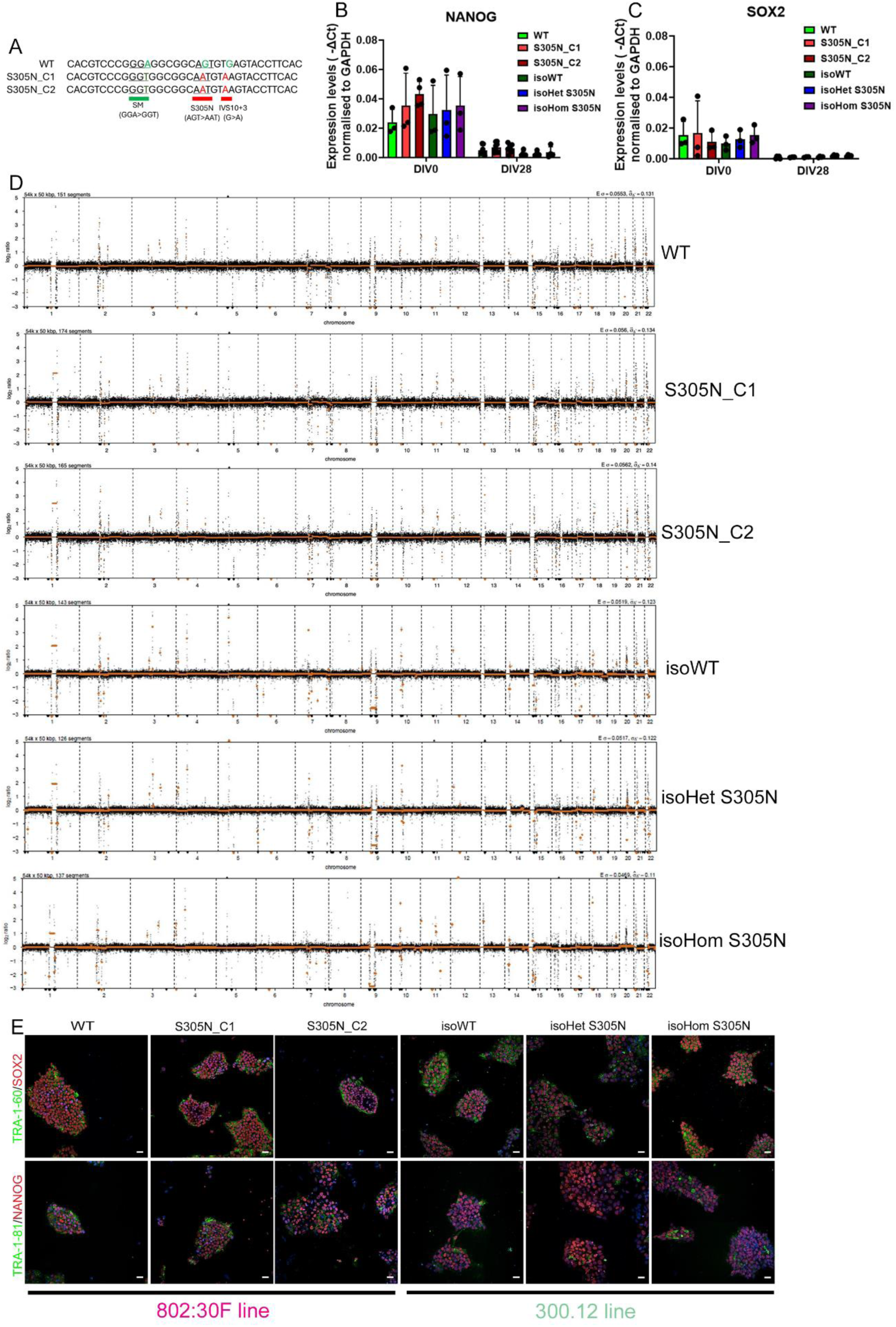
Characterisation of iPSCs with S305N tau mutations related to Fig. 1. (A) Schematic representation of the inserted mutations in the 802:30F iPSC line used to generate the S305N_C1 and S305N_C2 clones. (B–C) mRNA expression levels of the pluripotency markers NANOG (B) and SOX2 (C) in all lines at DIV0 (iPSC stage) and DIV28 (i^3^N neurons). Data are presented as mean ± SD, n = 3–4 independent neuronal differentiations. (D) Low-coverage whole-genome sequencing revealed no chromosomal abnormalities in any of the lines following hNGN2 insertion. (E) Immunostaining for pluripotency markers TRA-1-60/SOX2 and TRA-1-81/NANOG confirmed the maintenance of pluripotency in all iPSC lines after hNGN2 integration.

**Fig. S2.**
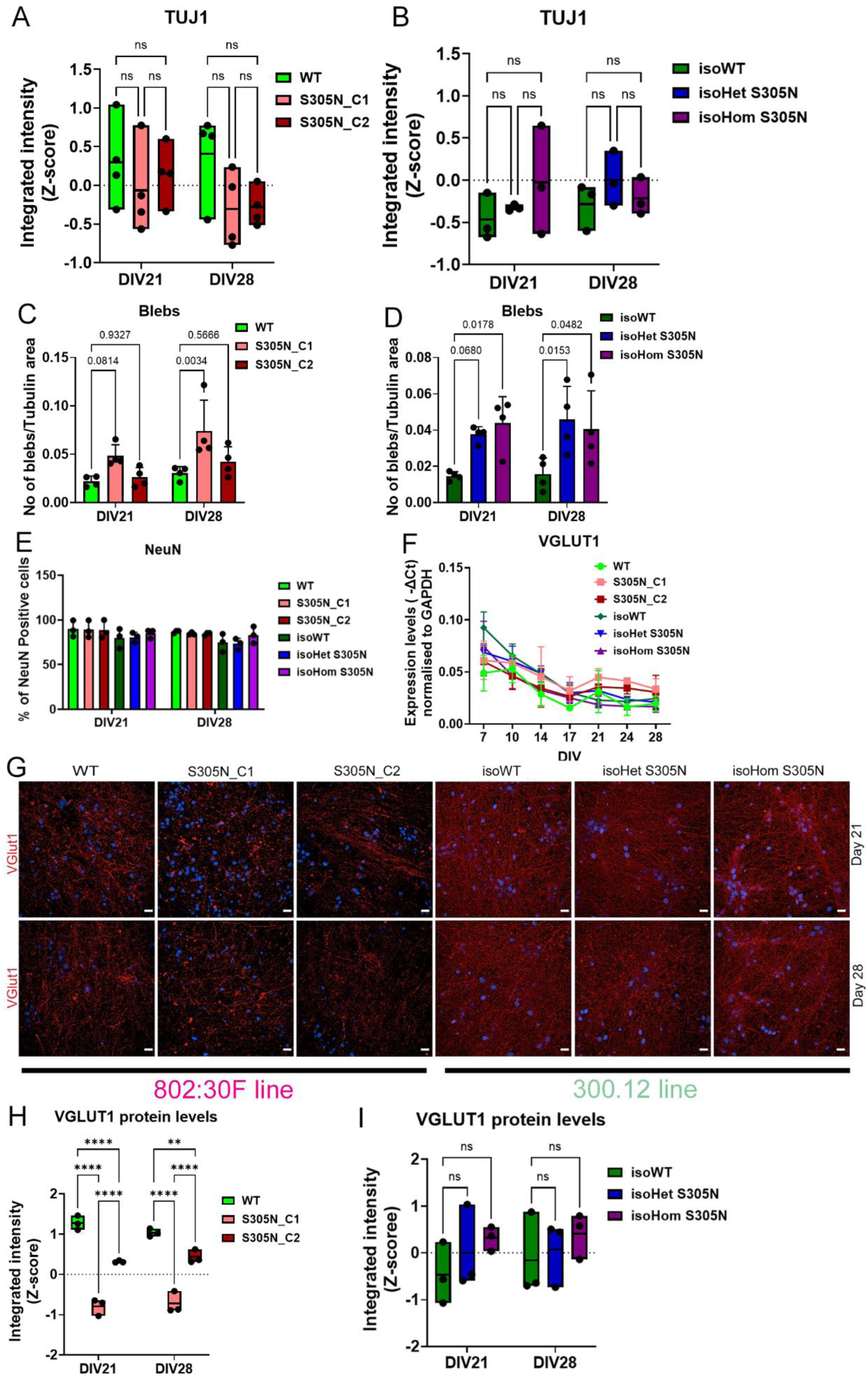
Characterising S305N i^3^N neurons related to Fig. 1. (A) Expression levels of TUJ1 in WT and S305N_C1 and C2 neurons at DIV21 and DIV28. (B) Expression levels of TUJ1 in isoWT, isoHet, and isoHom S305N neurons at DIV21 and DIV28. (C) Quantification of the number of blebs per TUJ1-positive area in WT and S305N_C1 and C2 neurons at DIV21 and DIV28. (D) Quantification of blebs per TUJ1-positive area in isoWT, isoHet, and isoHom S305N neurons at DIV21 and DIV28. (E) Percentage of NeuN-positive neurons in all lines at DIV21 and DIV28. (F) VGLUT1 mRNA levels across all lines over time. (G) Representative images of VGLUT1 staining in all neuronal lines at DIV21 and DIV28 (cells fixed with 4% PFA). (H) Quantification of VGLUT1 expression in WT and S305N_C1 and C2 neurons at DIV21 and DIV28. (I) Quantification of VGLUT1 expression in isoWT, isoHet, and isoHom S305N neurons at DIV21 and DIV28. Statistical analysis for all panels was performed using two-way ANOVA followed by Tukey’s post-hoc test. Data are presented as mean ± SD. Sample sizes: n = 3–5 independent neuronal differentiations. Significance: **p < 0.01 ****p < 0.0001.

**Fig. S3.**
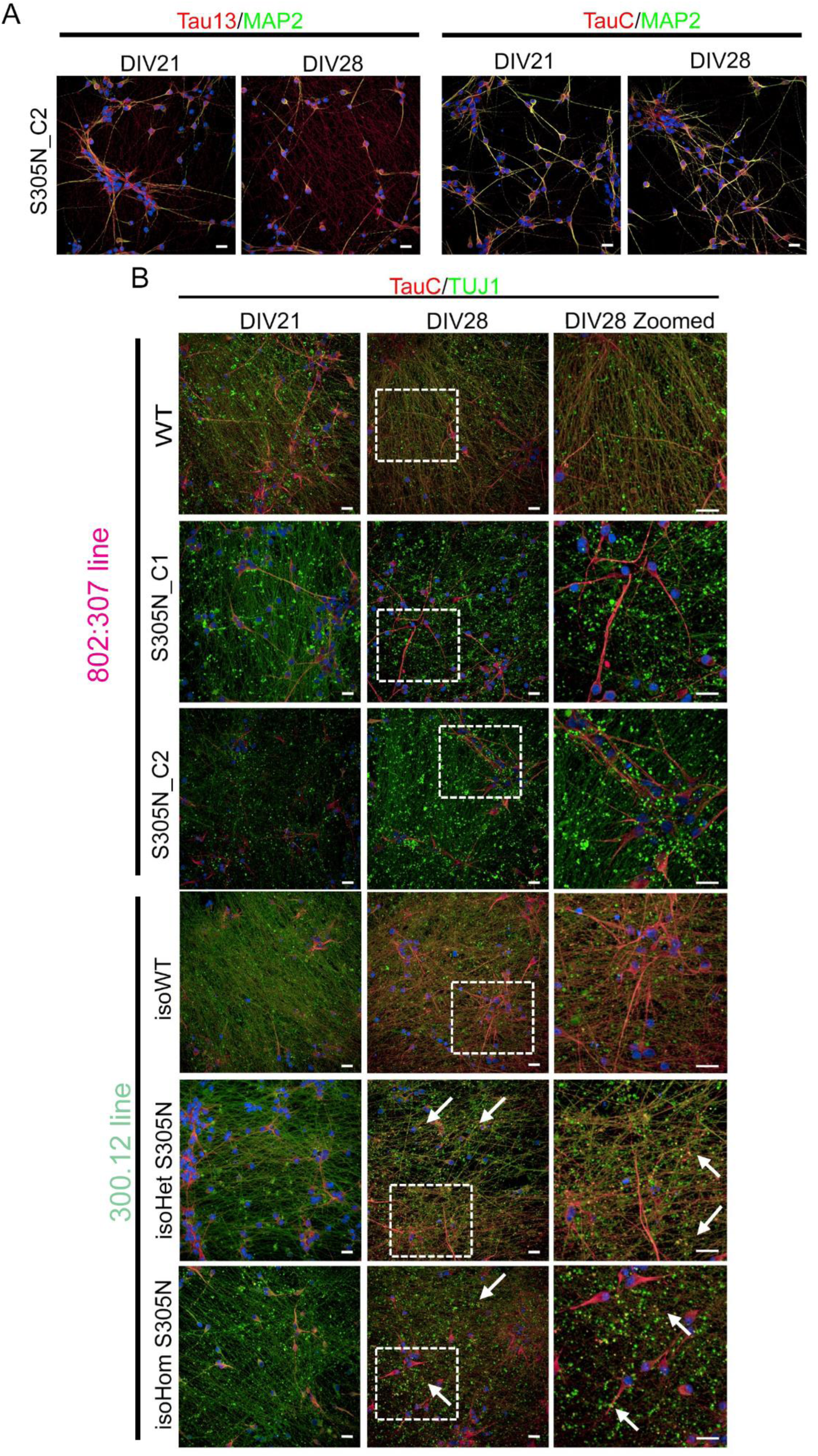
Distribution of total tau in S305N i^3^N neurons related to Fig. 2. (A) Representative images of DIV21 and DIV28 S305N_C2 i^3^N neurons double-labelled with MAP2 and total tau antibodies: Tau13 (left) and TauC (right). (B) Representative images showing TauC co-stained with TUJ1 across all neuronal lines. The enlarged image highlights colocalization of TauC-positive blebs with TUJ1-positive axonal blebs.

**Fig. S4.**
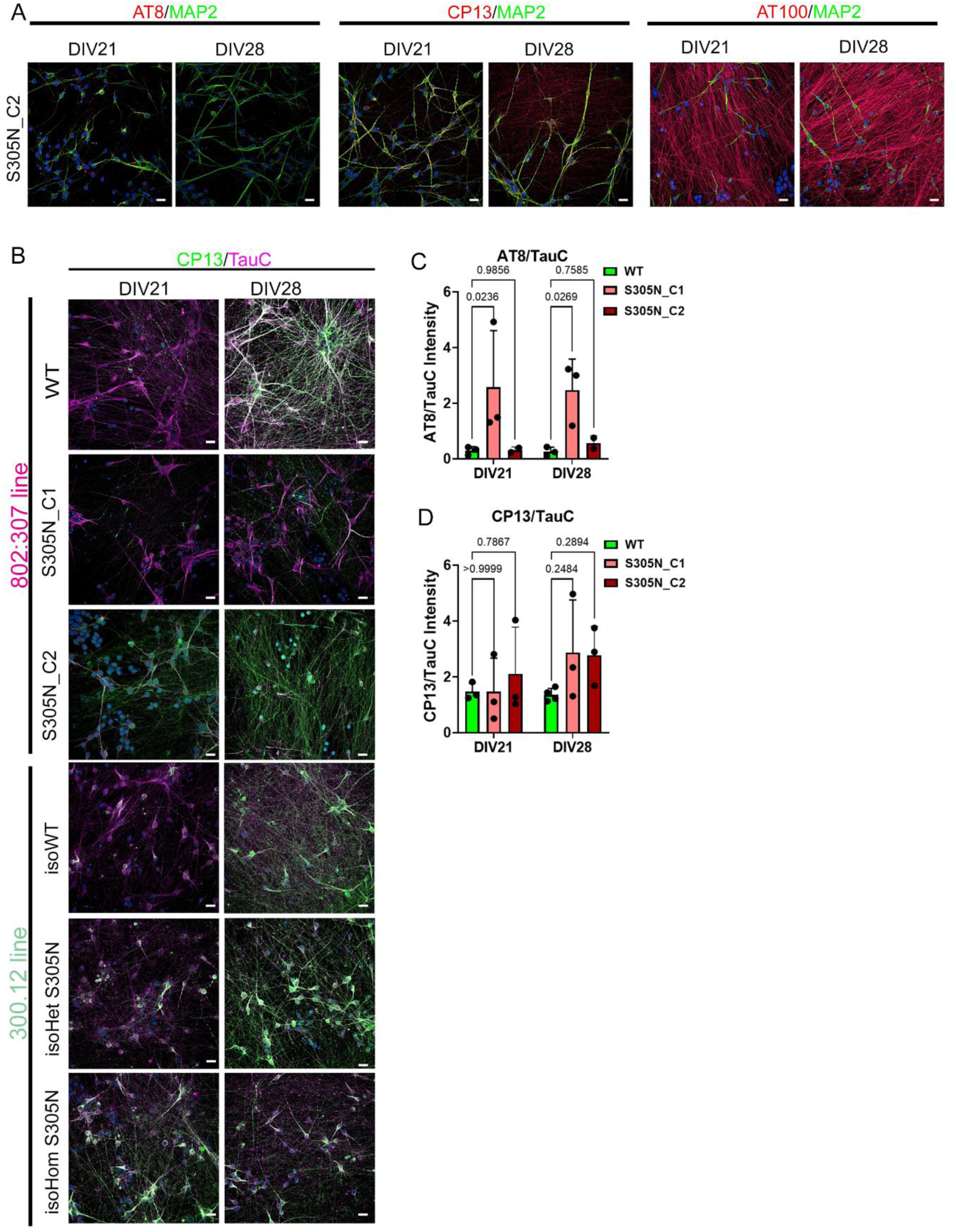
Distribution of phosphorylated tau in S305N i^3^N neurons related to Fig. 3. (A) Representative images of DIV21 and DIV28 S305N_C2 i^3^N neurons double-labelled with MAP2 and phosphorylated tau antibodies: AT8 (left), CP13 (middle), and AT100 (right). (B) Representative images showing co-staining of CP13 and TauC across all neuronal lines. (C) Quantification of AT8 immunostaining normalized to TauC in WT and S305N_C1 and C2 neurons at DIV21 and DIV28. Statistical analysis was performed using two-way ANOVA followed by Tukey’s post-hoc test (mean ± SD, n = 2–3 independent neuronal differentiations).(D) Quantification of CP13 immunostaining normalized to TauC in WT and S305N_C1 neurons at DIV21 and DIV28. Statistical analysis was performed using two-way ANOVA followed by Tukey’s post-hoc test (mean ± SD, n = 3–4).

**Fig. S5.**
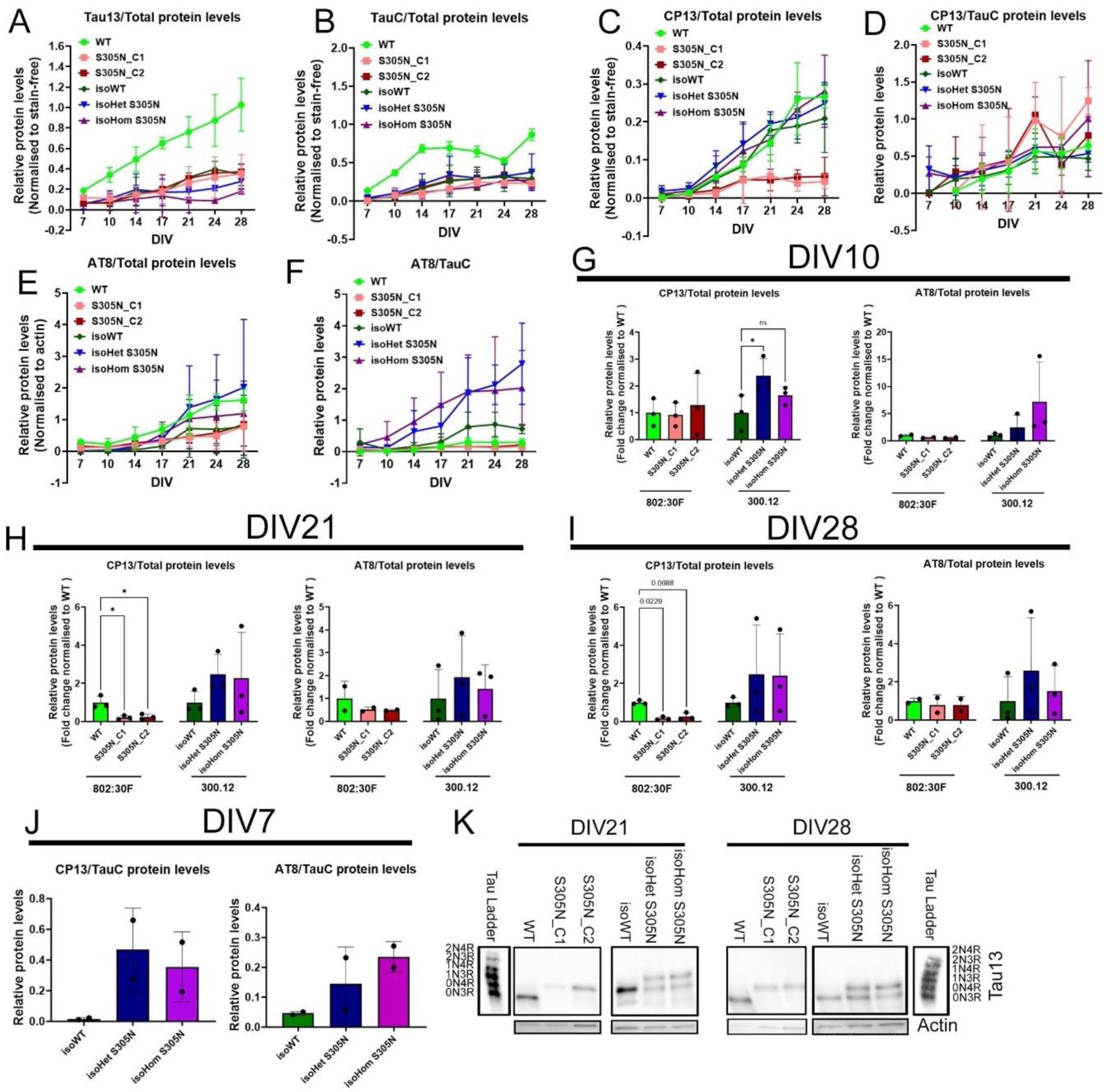
Expression levels of total and phosphorylated tau, related to Fig 4, over time. (A–B) Protein levels of total tau detected using Tau13 (A) and TauC (B) antibodies, normalized to total protein (stain-free imaging), across all neuronal lines over time. (C–D) Expression of phosphorylated tau detected using CP13, normalized to total protein (C) and to TauC (D), across all neuronal lines. (E–F) Expression of phosphorylated tau detected using AT8, normalized to total protein (E) and to TauC (F), across all neuronal lines. (G–I) Protein levels of CP13 (left) and AT8 (right) at DIV10 (G), DIV21 (H), and DIV28 (I), normalized to total protein (stain-free imaging) in all neuronal lines. (J) Protein levels of CP13 (left) and AT8 (right) normalized to TauC at DIV7 in isoWT, isoHet, and isoHom S305N neurons. (K) Representative immunoblots from the dephosphorylation assay for tau in all neuronal lines at DIV21 and DIV28. Tau13 was used to detect different tau isoforms. Statistical analysis was performed using repeated measures (RM) one-way ANOVA with Dunnett’s post-hoc test to compare S305N_C1 and C2 vs WT, and isoHet/isoHom S305N vs isoWT. These tests were selected due to variability in immunoblot replicates. Data are presented as mean ± SD; *, p<0.05; n = 2-3 independent neuronal differentiations.

**Fig. S6.**
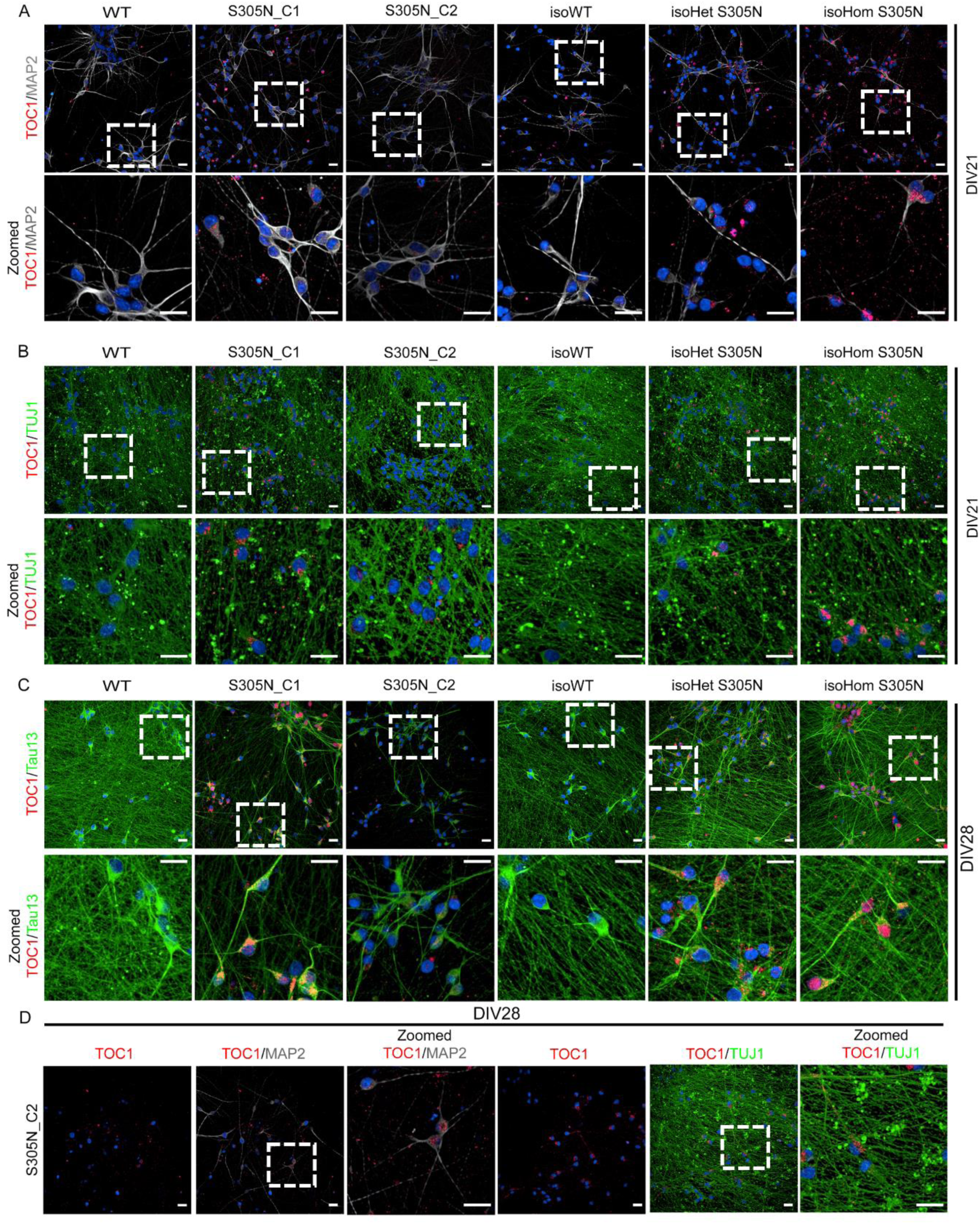
TOC1 expression at DIV21 and colocalization with Tau13 in S305N i^3^N neurons related to Fig. 5. (A) Representative images of TOC1 and MAP2 co-labelling at DIV21 for all neuronal lines. (B) Representative images of TOC1 and TUJ1 co-staining at DIV21 for all neuronal lines. (C) Representative images of TOC1 and Tau13 co-staining at DIV28 for all neuronal lines. (D) Representative images of S305N_C2 DIV28 i^3^N neurons stained with either TOC1 and MAP2 or TOC1 and TUJ1.

**Fig. S7.**
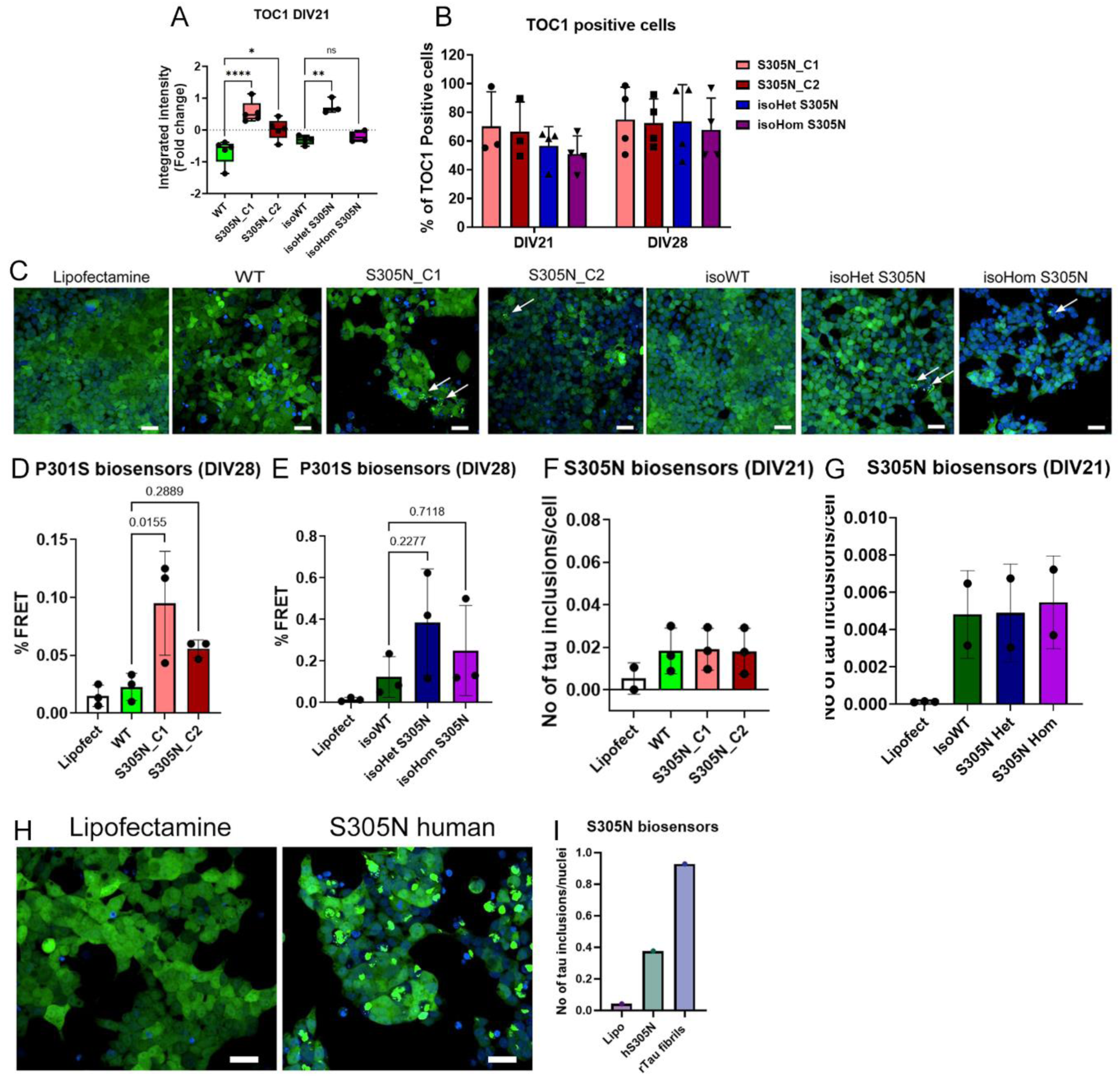
i^3^N neurons with a S305N mutation form endogenous seed-competent tau related to Fig. 5. (A) Quantification of TOC1 signal from DIV21 i^3^N neurons. Statistical analysis was performed using one-way ANOVA followed by Tukey’s post-hoc test to compare mutant lines with their respective isogenic controls (mean ± SD, n = 4–5 independent neuronal differentiations). (B) Percentage of TOC1-positive neurons in i^3^N cultures with S305N mutations. (C) Representative images showing tau seeding activity in S305N biosensors seeded with DIV28 i^3^N neuron lysates from all lines. (D–E) Quantification of seeding activity at DIV28 in P301S biosensors. Statistical analysis was performed using one-way ANOVA followed by Tukey’s post-hoc test (mean ± SD, n = 3 independent neuronal differentiations). (F–G) Quantification of seeding activity at DIV21 in S305N biosensors. Statistical analysis was performed using one-way ANOVA followed by Tukey’s post-hoc test (mean ± SD, n = 2–3 independent neuronal differentiations). (H) Representative images showing tau seeding activity in S305N biosensors seeded with human S305N brain tissue. (I) Quantification of seeding activity from 3 µg of frozen human S305N brain tissue using the S305N biosensors.

**Fig. S8.**
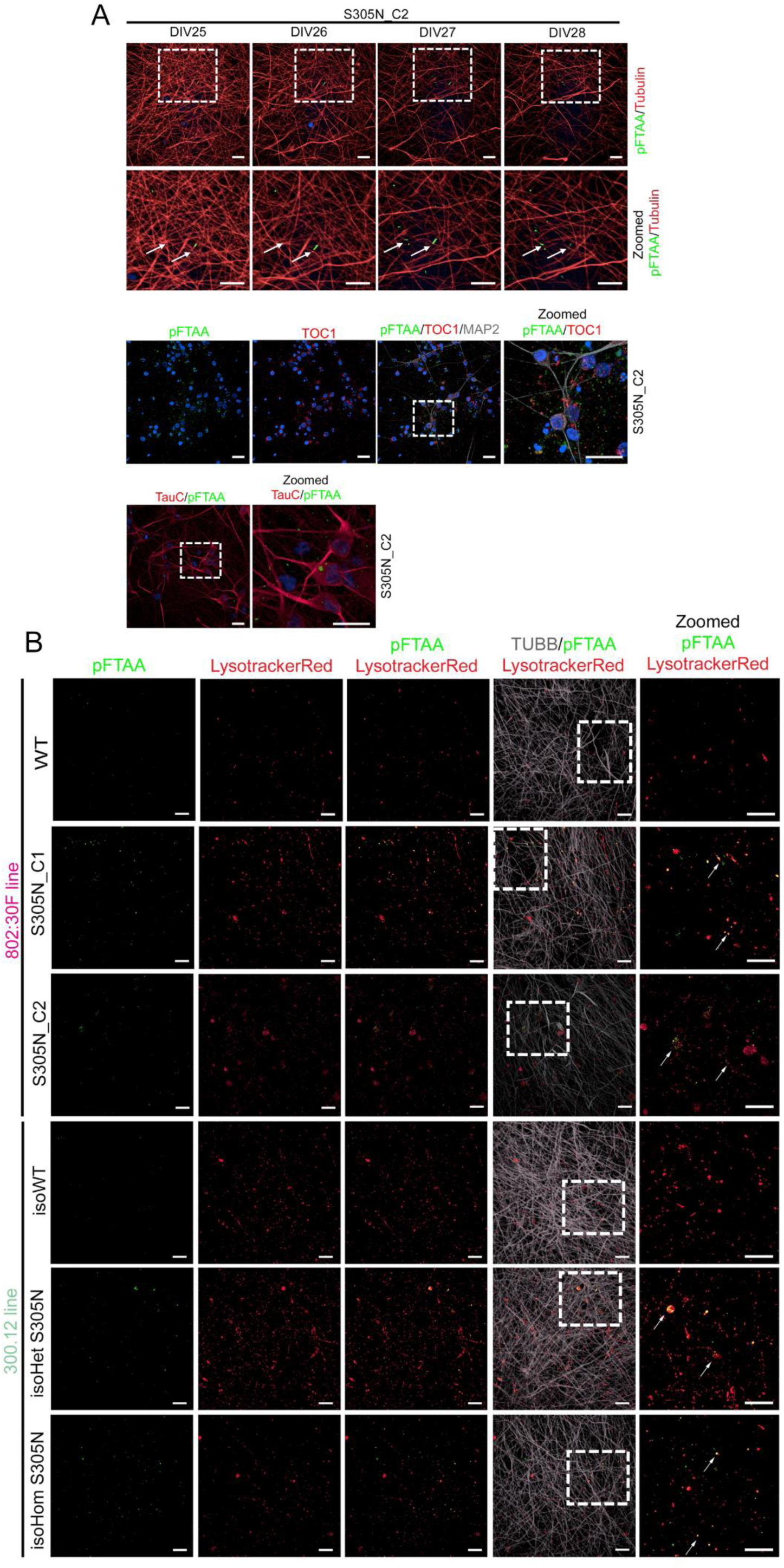
pFTAA colocalizes with LysotrackerRed in i^3^N neurons, related to Fig. 6. (A) Representative images of S305N_C2 i^3^N neurons stained live with pFTAA and Tubulin over time (top panel), co-stained with TOC1 and MAP2 (middle panel), and TauC (bottom panel). (B) Representative images of all neuronal lines stained with pFTAA, Lysotracker Red, and Tubulin (TUBB) at DIV28.

## Notes

### Competing Interest Statement

The authors have declared no competing interest.

### Summary of Updates

This version of the manuscript has been revised to include data from the second S305N_C2 clone for all experiments and add extra data on the seeding competent tau.

